# Metabolic feedback during bacterial fermentation is a motility brake

**DOI:** 10.64898/2026.04.18.717966

**Authors:** Lucas Le Nagard, Jana Schwarz-Linek, Ekaterina Krasnopeeva, Carine Douarche, Jochen Arlt, Angela Dawson, Vincent A. Martinez, Wilson C.K. Poon, Teuta Pilizota

**Affiliations:** Scottish Universities Physics Alliances and School of Physics and Astronomy, University of Edinburgh, James Clerk Maxwell Building, Peter Guthrie Tait Road, Edinburgh EH9 3FD, UK; School of Biological Sciences, King’s Buildings, EH9 3FF, University of Edinburgh, Edinburgh, UK; Department of Physics, Cavendish Laboratory, University of Cambridge, Cambridge, CB3 0HE, UK; Institute of Science and Technology Austria, Am Campus 1, 3400 Klosterneuburg, Austria; Université Paris-Saclay, CNRS, FAST, 91405 Orsay, France

**Keywords:** Physiology, Motility, Homeostasis, Proton motive force, Fermentation

## Abstract

We study an unexpectedly fast decay of motility in dense suspensions of *Escherichia coli* bacteria supplied with excess glucose under anaerobic conditions. The decrease in swimming speed occurs on a timescale inversely proportional to the cell concentration, and is associated with the secretion of organic acids by the bacteria. We show that the decay is driven by the progressive accumulation of non-ionised organic acids in the medium, and develop a chemical kinetic model that successfully predicts the swimming speed variations over a range of conditions in the presence of these acids. We further measure the internal pH of *E. coli* cells exposed to organic acids, and find that the speed decay coincides with sharp declines in internal pH and metabolic rate. Our findings identify an additional layer of motility control that can arise in complex environments even when motility genes are expressed and energy sources are abundant. This mechanism is likely relevant for understanding bacterial motility in habitats such as the human gut, where high densities of bacteria and organic acids are common.

---

Bacterial taxis allows cells to bias their movement toward favorable environments and away from harmful conditions [1– 3]. The observation of higher levels of expression of genes related to motility in poor nutrient conditions originally led to the conclusion that motility is a foraging response triggered by the scarcity of nutrients [4–6]. However, we now know that taxis also accelerates the expansion of bacterial colonies growing under nutrient-replete conditions [7]. The inverse correlation between the expression of motility genes and the growth rate was also recently shown to minimise the cost of keeping an entire bacterial population motile under various growth conditions [8, 9], further indicating that motility is not just a foraging response. This new paradigm expands the range of environments in which motility may be relevant. For *E. coli*, these include the urinary and gastrointestinal (GI) tracts of humans and other warm-blooded animals [10–12], soil and sediments [13, 14], and even the surface of algae and periphytons [13].

Exactly how motility contributes to the life of *E. coli* in these diverse environments is not known, but previous work suggested niche areas in the human GI tract where it is important, such as the oxygen-rich vicinity of the epithelial tissue protected by a layer of mucus [15]. The rest of the human large intestine, which harbours up to 10^11^ bacteria/mL [16], is anerobic, with the oxygen that diffuses through the intestinal epithelium being quickly consumed by the gut microbiome [17, 18]. Because *E. coli* cells can remain motile under anaerobic conditions [19], there is no *a priori* reason why they may not be able to swim in the intestinal lumen too. Under anaerobic conditions, *E. coli* and many other microbes produce short-chain fatty acids (SCFAs) by fermenting available sugars [20, 21]. At high concentrations, SCFAs are detrimental to microbes and as so are widely used as food preservatives. They negatively affect bacterial physiology by disrupting internal pH homeostasis [22–24], and slowing down growth [25–27]. Effects on motility have also been reported [22, 24, 28]. In particular, Minamino *et*.*al*. exposed dilute suspensions of *E. coli* to externally supplied SCFAs under aerobic conditions and found that SCFA-induced cytoplasmic acidification can slow and even prevent the rotation of bacterial flagellar motors [28]. However, it is unknown whether the endogenous production of SCFAs by bacteria can influence their own motility; if so, this could act as an additional regulatory layer.

Here, we first sought to determine whether such regulation occurs and, if so, to quantify its effect. For this purpose, we studied suspensions of motile *E. coli* cells in media where they cannot grow but can ferment glucose to generate a proton motive force and power their motility. By varying the buffering strength of the medium and cell densities, we observed that at about 10% of the total bacterial concentration found in the human colon, cells quickly produce enough SCFA to self-poison and become non-motile. Motility is recovered after SCFAs are removed, indicating a reversible effect on cellular physiology without damaging the flagella. We capture our results in a quantitative model that successfully links the effect of SCFA on motility with the concentration of protonated organic acids. Our findings suggest that the motility of *E. coli* can be important in the GI tract only if protonated organic acids concentrations remain low. Thus, given the low buffering capacity of the human GI tract content (20-40 mmol H^+^*/*L*/*ΔpH in the ascending colon [29], and ≲ 5mmol H^+^*/*L*/*ΔpH in the small intestine [30]), comparable to our experiments, this would require a fine balance between the flow of content, SCFA production by the microbiome, and their absorption and fresh buffer secretion by the host.

## Results

### In anaerobic environments and at high cell density, motile *E. coli* suspensions rapidly become non-motile

To determine whether self-production of SCFAs can influence the motility of *E. coli*, we conducted the study in nitrogen-free potassium phosphate buffer (Motility Buffer - MB) supplemented with 20 mM glucose (*Methods*). Working with non-growing bacteria ensured we could control the cell density while limiting the cells’ ability to adapt to environmental changes, and we know that the cells can remain motile for prolonged periods of time in this medium [31]. We measured swimming speeds with differential dynamic microscopy (DDM), a high-throughput method that can characterise the motility of thousands of cells simultaneously [32, 33]. DDM was performed in sealed glass capillaries (*Methods*), in which dissolved oxygen was depleted at a time that depends on the cell concentration [31].

We started with ≈ 2.5 × 10^10^ cellsmL^−1^, OD = 15. Extrapolating from previous work at considerably lower cell densities and glucose concentrations (OD ≲ 1 and ≲ 1mM respectively) [31], we expect these cells to exhaust dissolved oxygen in ≲ 1 min, but continue to swim at a constant speed for ≈ 100 min until they consume glucose [31]. Instead, the initially constant swimming speed first increased between *t* ≈ 8 min and *t* ≈ 12 min due to the emergence of collective motion in the suspension, Fig. SI1, and then dropped sharply to zero between *t* ≈ 13 min and *t* ≈ 25 min, Fig. 1(a). Similarly, abrupt cessation of motility was observed at lower cell densities, where there was no collective motion at the onset of the experiment, Fig. 1(b) and Fig. SI2. The time required for the speed to decrease by 50% from its maximal value is inversely proportional to the cell density, Fig. 1(b) (inset), suggesting that this timescale is set by glucose consumption and/or the production of compounds by the bacteria that stop motility.

**Fig. 1.**
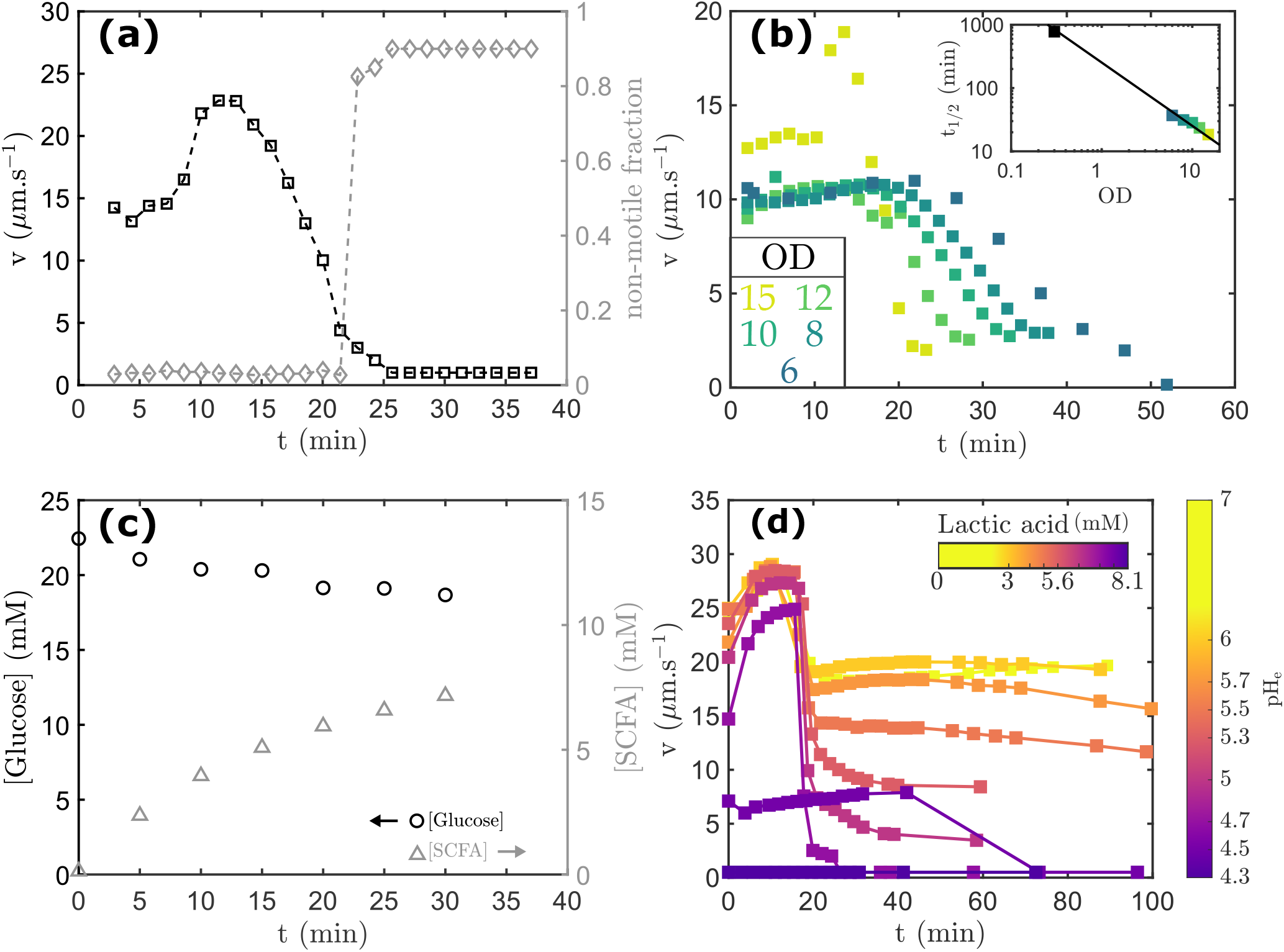
Dense, anaerobic suspensions of *E. coli* cells lose motility in glucose-supplemented Motility Buffer. **(a)** Average swimming speed (black squares) and non-motile fraction (grey diamonds) of a dense suspension (OD=15) under anaerobic conditions, displaying collective motion at *t* ≈ 10 min and a rapid decay of swimming speed for *t* ≳ 15 min. **(b)** Swimming speed as a function of time for *E. coli* suspensions of varying cell densities. Inset: *t*_1*/*2_, the time required for the speed to decay by 50%, plotted as a function of cell density. The colors correspond to the main panel, with an added black square corresponding to OD=0.3 (see Fig. SI2). Black fit line: *f* (*x*) = *a/x* with *a* = 253 min OD. **(c)** Glucose (black circles) and SCFA (gray triangles) concentrations in the Motility Buffer as a function of time, for a dense suspension from (a) (OD=15). **(d)** Swimming speeds of dilute *E. coli* suspensions (OD = 0.3) in Motility Buffer + 20 mM glucose supplemented with varying lactic acid concentrations, from yellow to purple: 0 mM & pH_e_ = 7.0, 3 mM & pH_e_ = 6.0, 4.2 mM & pH_e_ = 5.7, 5.1 mM & pH_e_ = 5.5, 5.6 mM & pH_e_ = 5.3, 6.2 mM & pH_e_ = 5.0, 6.9 mM & pH_e_ = 4.7, 7.3 mM & pH_e_ = 4.5 and 8.1 mM & pH_e_ = 4.3. The initial fast speed regime (*t* ≲ 20 min for most curves, *t* ≲ 40 min for 7.3 mM lactic acid) corresponds to aerobic conditions, before cells have depleted the dissolved oxygen. Lines are provided as guides to the eye.

To distinguish between these two possibilities, we combined nuclear magnetic resonance (NMR) with ion chromatography to identify and quantify SCFAs in a dense suspension (OD = 15) as a function of time (*Methods*, Fig. SI3), and measured the glucose concentration at the same time points (*Methods*), Fig. 1(c). Glucose remained abundant throughout the experiment, excluding substrate depletion as a cause of the speed decay. On the other hand, the SCFA concentration in the medium increased non-linearly with time, Fig. 1(c), pointing to this as a potential cause. We further found that a strain that does not tumble exhibits the same motility decay as the wild type, and that a non-motile mutant exhibits the same SCFA production pattern (*Methods*, Fig. SI4). Thus, the speed decay is not related to negative taxis and tumbling, and variations in metabolic activity are independent of motility.

To assess whether the speed decay was directly related to SCFAs, we prepared dilute cell suspensions (OD = 0.3) supplemented with varying concentrations of the most abundant SCFA detected in Fig. 1(c), lactic acid (see *Methods* and Table SI1 for SCFA quantification). Dissolved oxygen was not immediately depleted at this low cell density, so that we observe two speed regimes, Fig. 1(d): an aerobic one characterised by fast speed, followed by a slower anaerobic one. The addition of lactic acid caused a concentration-dependent decay of swimming speed in both regimes, although higher concentrations were required to produce a significant decay under aerobic conditions. Taken together, these results suggest that dense suspensions rapidly become non-motile under anaerobic conditions due to bacterial production of SCFAs.

### Extracellular pH_e_ and SCFA concentration do not independently govern the motility decay

Fermentation products can impair bacterial processes such as swimming, either by acting as toxins or by lowering the pH of the medium. To distinguish between these possibilities, we repeated the motility measurements at the same cell density (OD = 15) and glucose concentration (20 mM) in a ten times more concentrated phosphate buffer (10× MB, see *Methods*) to resist SCFA-induced pH_e_ variations^1^. Motility was maintained for ≈ 100 min in this buffer, Fig. 2(a), a significant increase compared to 1× MB + 20 mM glucose. However, the speed again continuously decreased after reaching a maximum at *t* ≈ 30 min. The low initial speed in 10× MB + 20 mM glucose compared to 1× MB + 20 mM glucose was caused by the larger osmolarity (≈ +235 mOsm compared to 1× MB + 20 mM glucose), and the initial speed increase corresponds to the slow recovery from the osmotic shock [31, 34, 35]. Here, we did not observe signature of collective motion. Furthermore, increasing the amount of glucose did not affect the motility decay, ruling out once again the possibility that it was caused by glucose depletion, Fig. SI5.

**Fig. 2.**
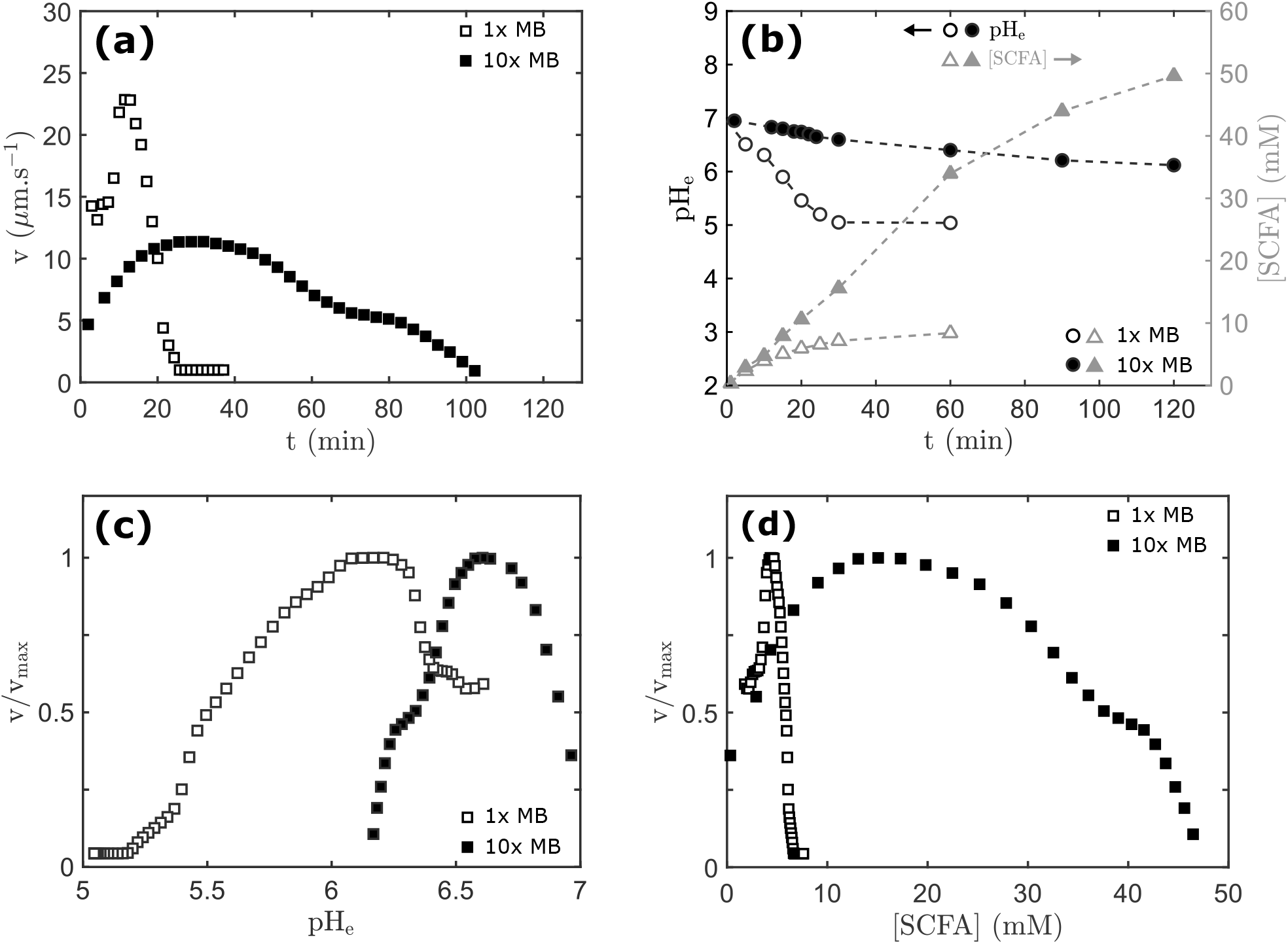
pH_e_ and [SCFA] do not independently dictate the swimming speed decay. **(a)** Average swimming speed of dense suspensions (OD=15) as a function of time in 1× MB + 20 mM glucose (open squares) and 10× MB + 20 mM glucose (filled squares), showing that cells remain motile for a longer period of time in 10× MB + 20 mM glucose. **(b)** pH_e_ (circles) and SCFA concentration (triangles) as a function of time for dense suspensions (OD=15) in 1× MB + 20 mM glucose (open markers) and 10× MB + 20 mM glucose (filled markers). SCFA concentrations reported for 10× MB at *t* = 90 min and *t* = 120 min were computed from glucose measurements (see *Methods*). **(c)** Normalised swimming speed as a function of pH_e_ and **(d)** SCFA concentration for dense suspensions in 1× MB + 20 mM glucose (open squares) and 10× MB + 20 mM glucose (filled squares).

pH_e_ decreased in both media, but more slowly and to a smaller extent in 10× MB + 20 mM glucose than 1× MB + 20 mM glucose, Fig. 2(b), which is reminiscent of the slower speed decay observed in the stronger buffer. The SCFA concentration was similar in both media during the initial ≈ 10 min, Fig. 2(b). Then, the SCFA concentration increased more slowly in 1× MB + 20 mM glucose, while it continued to increase at the same rate in 10× MB + 20 mM glucose until *t* ≈ 60 min. The onset of speed decay was observed at *t* ≈ 10 min in 1× MB + 20 mM glucose, and the middle of the decay at *t* ≈ 60 min in 10× MB + 20 mM glucose. Thus, the speed decay coincides with a slowdown of the metabolic activity.

To link these measurements with the speeds measured in each buffer, we interpolated pH_e_, SCFA concentration, and swimming speeds at the same time points, Fig SI6. We found that pH_e_ did not dictate the speed changes. For example, cells were non-motile in 10× MB + 20 mM glucose at the final pH_e_ of ≈ 6.1, while they were highly motile at the same pH_e_ in 1× MB + 20 mM glucose, and still swam at ≈ 50% of their maximum speed at pH_e_ = 5.5, Fig. 2(c). The concentration of SCFA in the medium also did not govern the speed changes in both buffers, Fig. 2(d). For example, ≈ 37 mM SCFA were required to induce a 50% speed decay in 10× MB + 20 mM glucose, compared to only ≈ 6 mM in 1× MB + 20 mM glucose.

### The accumulation of protonated SCFA drives the motility decay

Since carbon source availability, pH_e_, total [SCFA], and the duration of the experiment could not independently account for the motility decay, we hypothesized that the main driver was the accumulation of their uncharged form, SCFAH. Unlike SCFA^−^ anions, SCFAHs readily diffuse through lipid membranes [36, 37] and can therefore directly affect internal physiological processes, Fig. 3(a). If our hypothesis is correct, and assuming that intra- and extracellular SCFAH concentrations equilibrate rapidly so that [SCFAH]_e_ is a good proxy for [SCFAH]_i_, different speed measurements should all collapse onto the same ‘master curve’ when replotted against the [SCFAH] present in the buffers.

**Fig. 3.**
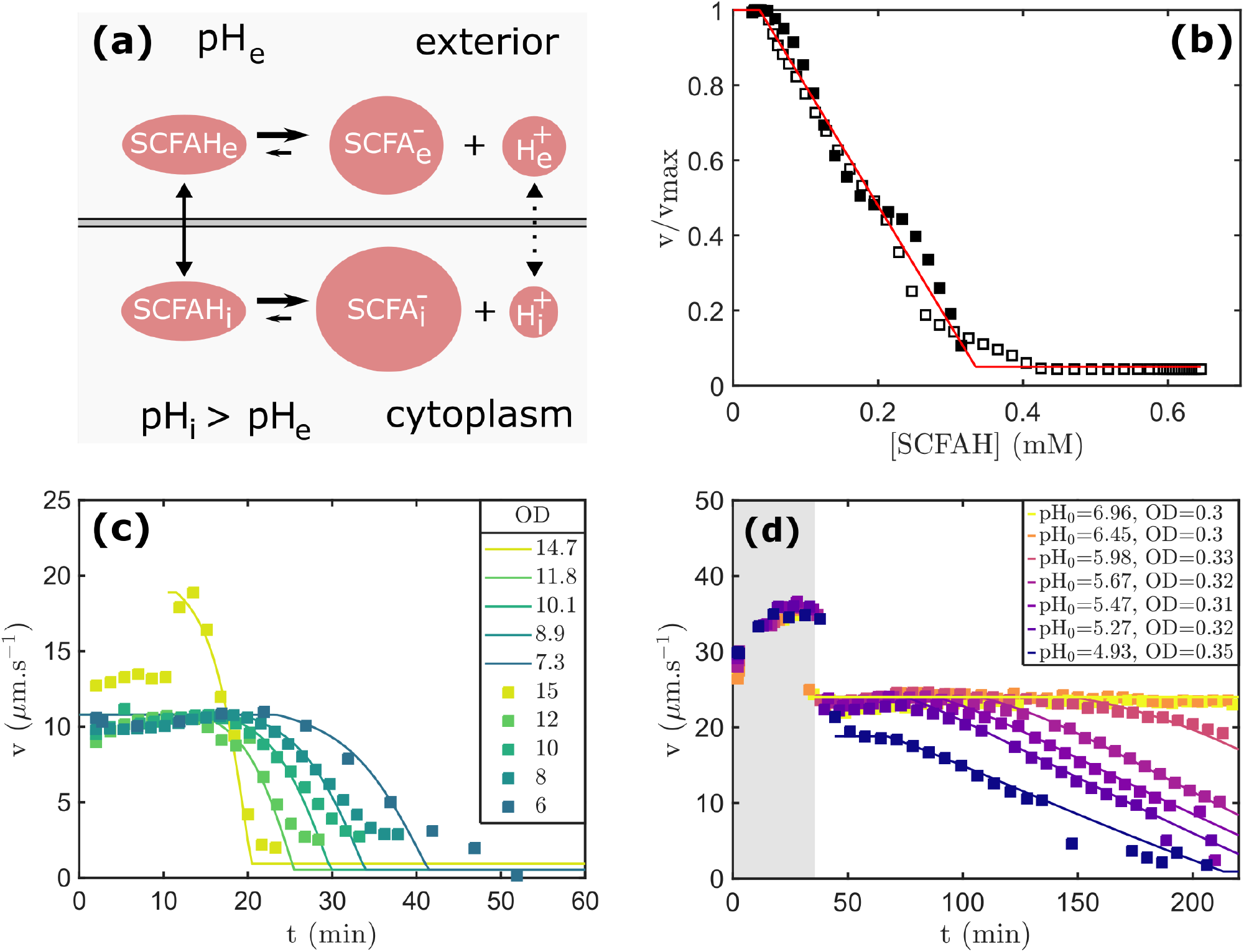
SCFAH govern the swimming speed decay. **(a)** Schematic of SCFA equilibration across the inner membrane of *E. coli*, where we assume the periplasmic space is part of the exterior environment. Protonation/deprotonation reactions are fast, such that SCFAH and SCFA^−^ are assumed to be in equilibrium both inside and outside the cell. Uncharged SCFAH diffuse across the membrane (solid transmembrane arrow), such that [SCFAH]_i_ ≈ [SCFAH]_e_, unlike deprotonated SCFA^−^ that do not. Protons can be rapidly transported across the membrane (dashed double arrow). **(b)** Normalised speeds display the same decay in 1× (open markers) and 10× (filled markers) MB + 20 mM glucose as a function of [SCFAH], indicating that [SCFAH] is the primary driver of the observed motility decay. Red line: fit to both curves with the function *f* (*x*) = min(1, max(*v*_∞_, (1 − *α*(*x* − *c*_A_)))) returning *v*_∞_ = 0.050, *α* = 3.20 mM^−1^, *c*_A_ = 0.037 mM. **(c)** Experimental (squares) and numerical (continuous lines) results obtained for bacterial suspensions with a range of ODs. The initial _e,0_ is 7.0 in all cases. **(d)** Experimental and numerical results obtained for low-density bacterial suspensions (OD = 0.3 in experiments) with a range of initial pH_e,0_. OD values in the numerics were manually fitted to best match the experimental results. Grey area: aerobic regime, during which no or negligible amounts of SCFA are produced.

To obtain [SCFAH] in our experiments, we calculate the concentration of uncharged monoprotic and diprotic acids in our mix at a given pH_e_ using

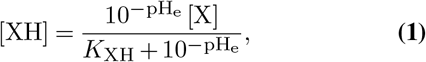

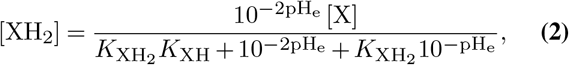

where [X] = [XH] +[X^−^] or [X] = [XH_2_] +[XH^−^] +[X^2−^] denote the concentrations of the protonated and deprotonated forms of monoprotic and diprotic acids, all at equilibrium. Here, *K* are their corresponding dissociation constants. These equations come from standard systems of equations for acid-base equilibria:

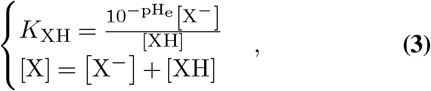

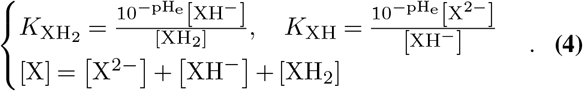

We use the measured pH_e_, Fig. 2(b), and dissociation constants from tabulated values (SI text 1 and Table SI2). We also need the total extracellular concentration, [X], of each acid to obtain the corresponding [XH] or [XH_2_] from Eq. 1 and Eq. 2. At several time points, we identified the fermentation products using NMR and quantified them using ion chromatography (*Methods*, Table SI1). The relative composition of the SCFA mixture remained the same at all times *t* ≥ 5 min. We therefore calculate [XH] and [XH_2_] using this time-invariant composition: 57.1% lactic acid, 21.7% formic acid, 14.6% acetic acid, and 6.6% succinic acid (*Methods*). We neglect ethanol, which is not an acid and was produced at concentrations that did not affect swimming, Fig. SI7. With all of these values in hand, we calculate the total [SCFAH](t), Fig. SI8.

Finally, after interpolating [SCFAH](t) in Fig. SI8 to match the time points at which swimming speeds are available, we plot the normalised swimming speed as a function of [SCFAH], Fig. 3(b). The data in both 1× and 10× MB + 20 mM glucose buffers collapse onto a master curve:

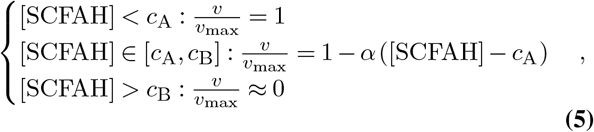

Here, *c*_A_ = 0.037 mM and *c*_B_ = 0.33 mM are the experimentally-determined maximum [SCFAH] that *E*. coli tolerate without affecting its swimming speed, and the [SCFAH] at which the swimming speed vanishes, respectively. These values hold for the particular SCFA mixture produced under our conditions. The parameter *α* = 3.2 mM^−1^ characterises the sharpness of the speed decay for [SCFAH]*> c*_A_.

### An empirical model predicts the SCFA-induced motility decay for various cell densities and pH_e_

Having demonstrated that [SCFAH] is the main driver of motility decay for dense suspensions in 1× and 10× MB + 20 mM glucose initially at pH_e_ = 7, we now test whether this is applicable to a wider range of bacterial densities and initial pH_e_, e.g. for the data presented in Fig. 1(b). To do so, we use Eq. 5 to predict the swimming speed from [SCFAH] for a given condition, and compare the calculated values with the experimental data. Similarly to above, [SCFAH] is calculated from [SCFA], pH_e_, *K*, and the relative composition of SCFA. We compute [SCFA] at each time step in the numerics using:

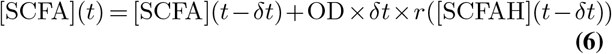

with *δt* = 0.5 min and *r* the SCFA production rate, expressed in mMmin^−1^ OD^−1^. *r*(*t*) was derived from measurements of [SCFA] in 1× MB + 20 mM glucose at OD=15. We further assumed that *r*(*t*) is a function of [SCFAH], and derived an empirical relationship for *r*([SCFAH]), Fig. SI9.

From [SCFA](*t*), we obtain the concentration of each SCFA, assuming that the relative composition of the SCFA mixture remains the same. We then calculate pH_e_ and [SCFAH] by solving a system of 14 equations that include charge balance, the dissociation equilibrium of each acid, conservation of matter, and self-ionization of water (see SI text 2 and Fig. SI10). We obtain the normalised swimming speed from [SCFAH] and Eq. 5, and finally multiply it by the maximum anaerobic swimming speed to obtain the absolute speed. We repeat this procedure for each time step to simulate the temporal evolution of the swimming speed (see SI text 3).

We first return to the data of Fig. 1(b), where we considered dense bacterial suspensions at various cell concentrations. The numerics capture the observed slowed build-up of [SCFAH] with decreasing bacterial concentrations, and consequently, the delayed motility crash, Fig 3(c). With the exception of OD = 6 (~ 20% discrepancy), experimental and model-predicted OD values differ by ≲ 10% during the initial decay phase (*v* ≳ 3 µms^−1^), consistent with the expected uncertainties in buffer preparation and bacterial dilution. Good agreement was also obtained on longer timescales in the anaerobic regime of an experiment conducted at OD=0.3, although the final stages of the decay were slower than predicted by the model, Fig. SI11.

We also wish to investigate the evolution of the anaerobic speed for a range of initial pH_e_ values. In particular, we predict that adjusting the initial pH_e_ without external addition of SCFA should initially have a minor effect on swimming speed compared to adding an SCFA. However, a lower pH_e_ should lead to a higher [SCFAH] for a given [SCFA]. From this, we expect the speed to decrease more rapidly as bacteria release SCFAs in suspensions prepared at a lower initial pH_e_. In line with this expectation, varying the initial pH_e_ (between 4.93 and 6.96, Table SI3) by modifying the phosphate buffer of 1× MB + 20 mM glucose, left the swimming speed of dilute bacterial suspensions (OD = 0.3) mostly unaffected in both aerobic and anaerobic conditions, Fig. 3(d). Our numerics again captured the dynamics of decay, confirming that [SCFAH] is the key parameter affecting bacterial physiology in our conditions. The experiments of Fig. 3(d) also reveal that pH_e_ begins to independently affect bacterial physiology under anaerobic conditions at pH_e_ = 4.93. Because it accounts for ≈ 20% reduction of the anaerobic speed, at that pH_e_ we adjusted *v*_max_ in the numerics, and found that the rest of the speed decay was again well captured by our model.

### SCFAH-induced disruption of pH_i_ homeostasis is the primary driver of the observed decay in swimming speed

Having shown that an increasing [SCFAH] in the medium is responsible for the collapse of bacterial motility during fermentation, we proceed to identify its physiological cause.

The flagellar motors of *E. coli* are powered by the proton motive force (PMF), to which the cells’ swimming speed is proportional in aqueous media [38–40]: *v* = *β* · PMF and PMF = Δ*ψ* − ln(10) 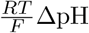, with *β* a (dimensional) constant, Δ*ψ* the membrane potential, ΔpH = pH_i_ − pH_e_, *R* the gas constant, *T* the temperature, and *F* the Faraday constant. Thus, one possibility is that the PMF decreases during exposure to SCFAH. Alternatively, because SCFAH readily diffuse through biological membranes, Fig. 3, their accumulation may negatively affect pH_i_ regulation [41] leading to acidification of the cytoplasm. Previous work reported that cytoplasmic acidification at nearly constant PMF during exposure to SCFA decreased the swimming speed of *E. coli* [28], thus a similar mechanism could be at play here. We first assessed the extent of pH_i_ disruption caused by fermentation products by exposing bacteria expressing the ratiometric pH sensor pHluorin (*Methods*, [42]) to cell-free, reoxygenated supernatant recovered from a dense suspension (OD = 15) after 30 min of fermentation (termed AB30 buffer, pH_e_ = 5.0). The initial pH_i_ = 7.9 *±* 0.6 in MB + 20 mM glucose quickly decreased to 6.0 *±* 0.3 upon exposure to AB30, Fig. 4(a), and recovered to pH_i_ = 7.8 *±* 0.5 when fresh buffer was flushed into the chamber at *t* = 17 min. A similar reversible effect was observed on the swimming speed of wild type bacteria, which decreased ≈ 40% upon exposure to AB30 under aerobic conditions and was fully restored after cells were transferred to fresh MB + 20 mM glucose, Fig. SI12.

**Fig. 4.**
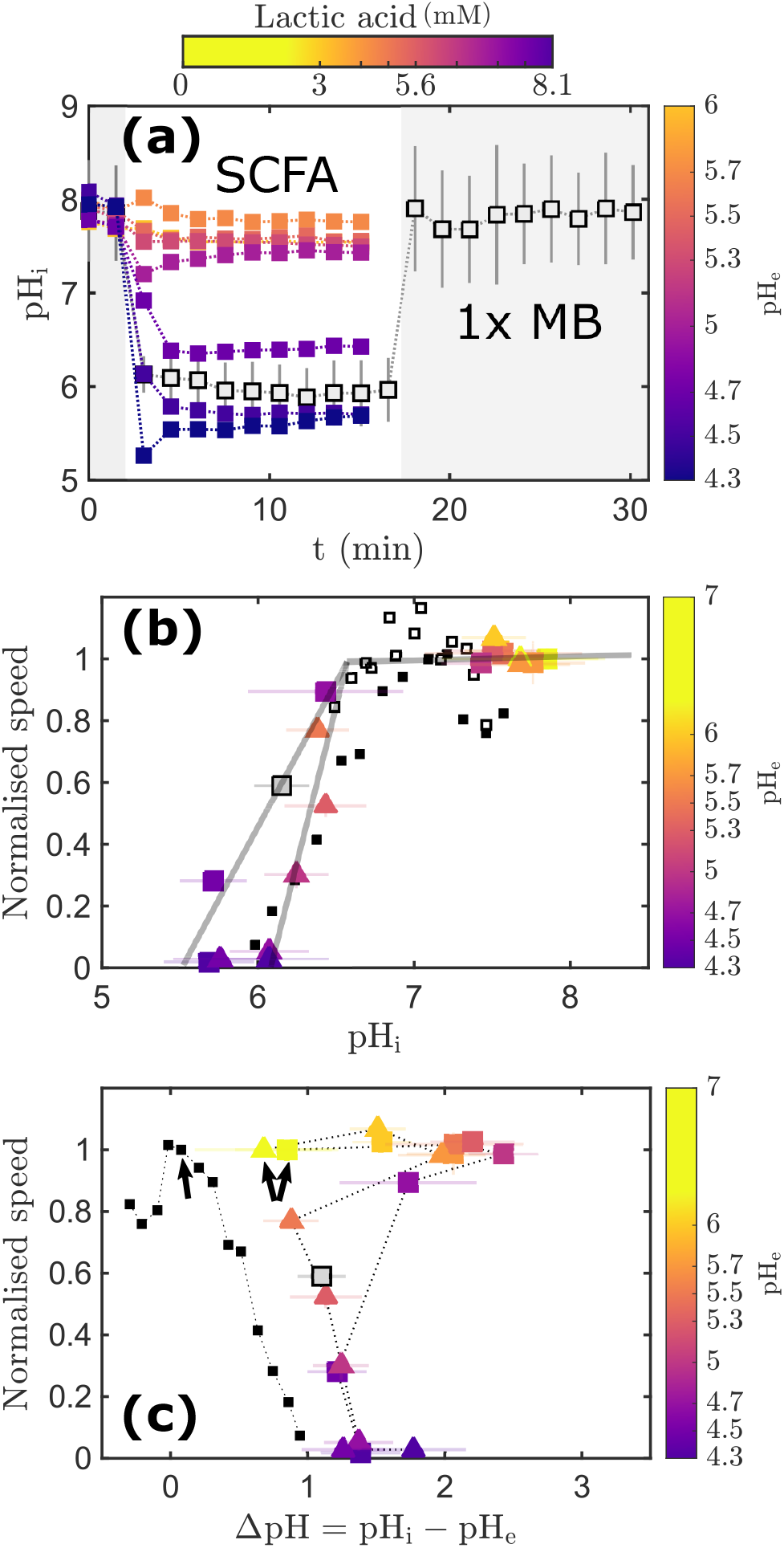
The swimming speed decay is accompanied by an SCFA-induced acidification of the cytoplasm. **(a)** pH_i_ of cells exposed to fermentation supernatant AB30 (grey squares), or to MB supplemented with lactic acid (colours as in Fig. 1(d)). Grey areas: 1× MB + 20 mM glucose, no SCFA. **(b)** Normalised speeds plotted against pH_i_, using pH_i_ data of panel (a), speed data of Fig. 1(d) and Fig. SI12, and data of Minamino *et al*. [28]. Squares and triangles depict aerobic and anaerobic results, respectively. Colours correspond to panel (a) and are our results obtained in 1× MB + 20 mM glucose + lactic acid, whereas filled and open black squares depict aerobic results obtained in MB (no glucose) + 10 mM lactate + 34 mM acetate, and in MB (no glucose) + 10 mM lactate (no glucose) by Minamino *et al*. [28], respectively. Lines are manually drawn to pass through the data points or the associated error bars, and thus highlight the difference between the aerobic and anaerobic conditions. **(c)** Normalised speeds plotted against ΔpH. In order to compare between studies, speeds in (b) and (c) are normalised by the speed measured at pH_e_=7.0. Dotted lines connect points in order of decreasing pH_e_, with points at pH_e_=7.0 indicated by arrows. Marker types and colors are as in (b). See *Methods* for the procedure used to plot the data of Ref. [28].

We further find that external addition of lactic acid also collapses pH_i_, Fig. 4(a) and Fig. SI13, and that this effect is correlated with that of lactic acid on swimming speed reported in Fig. 1(d). In aerobic conditions, speed and pH_i_ declines were first observed for an added concentration of lactic acid of 6.9 mM and pH_e_ = 4.7. Further increasing the concentration of lactic acid to 7.35 mM and 8.1 mM (pH_e_ = 4.5 and pH_e_ = 4.3) led to a dramatic decrease of aerobic speed to ≈ 7 µms^−1^ and 0 µms^−1^, accompanied by a further intracellular acidification to pH_i_ ≈ 5.8. Similar trends were recorded in anaerobic conditions (see *Methods*), Fig. SI14, Fig. 1(d); however, the speed decay began at a lower SCFA (and, therefore, SCFAH) concentration, with a ≈ 50% reduction in swimming speed already at 5.65 mM lactic acid and pH_e_ = 5.28.

Plotting the relative swimming speed variations against experimentally obtained pH_i_, Fig. 4(b) shows that the speed decay coincides with the loss of pH_i_ homeostasis (pH_i_ ≲ 6.7), in agreement with the results of Minamino *et al*. [28]. Furthermore, we consistently measured a large ΔpH even after complete decay of the swimming speed, Fig. 4(c). Together, these observations indicate that the main driver of SCFAH-induced speed decay is the effect of cytoplasmic acidification on flagellar motor rotation [28], which can be captured by defining *β* = *β* (pH_i_) in the relationship between speed and PMF, *v* = *β* (pH_i_) · PMF. However, we note that our aerobic and anaerobic speed decays do not collapse onto a single curve as a function of pH_i_, and the aerobic data of Ref. [28] overlap with our anaerobic rather than aerobic results. Furthermore, we also observed a decrease in metabolic activity as [SCFAH] increased, Fig. 2(b), which can cause a decrease in PMF. Given these observations, we conclude that small changes in PMF, and/or the influence of additional variables on *β* (pH_i_), have a minor contribution to the decline in swimming speed.

## Discussion

We conducted high-throughput motility measurements in dense bacterial suspensions fermenting glucose and observed sudden declines in motility and metabolic activity. We show that this physiological disruption is self-inflicted, and is governed by the accumulation of uncharged organic acids in the medium. Consistent with previous work [28,43], we find that the decay of swimming speed is primarily associated with a failure of pH_i_ homeostasis. Thus, accounting for the pH_i_ dependency in the relationship between the swimming speed and PMF, i.e., *v* = *β*(pH_i_) · PMF, explains the majority of the effect. Here we note that we know the linear relationship still holds at different pH_i_ from previous work [44]. The decrease in glucose consumption and SCFA production rates, concomitant with the speed decay in anaerobic conditions also agrees with previous reports that intracellular acidification inhibits glycolysis [45].

Different slopes recorded for the speed variations as a function of pH_i_ in aerobic and anaerobic conditions in Fig. 4(b), combined with the differences observed with previous results [28], suggest that there are additional determinants of the speed decay in our experiments. For example, the decrease of *β*(pH_i_) during cytoplasmic acidification may be compounded by a decrease in PMF caused by the metabolic slowdown. Alternatively, *β* may be a function of more than just pH_i_. In particular, if an acidic pH_i_ affects the motor by slowing the release of protons translocated through the motor into the cytoplasm [28], then the relative effect on motor speed may itself be speed-dependent, making *β*(pH_i_) a functional of the (unperturbed) speed, and the speed-PMF relationship non-linear. This and our recent report of saturation of the linear relationship between motor speed and PMF at high torque [40] suggest that future characterisation of the magnitude and dependencies of *β* should be fruitful for understanding the input-output relationship of the flagellar motor.

Because SCFAH can diffuse across cell membranes [36, 37], intracellular acidification during SCFA supplementation may arise from external SCFAH directly carrying protons into the cytoplasm, as long as [SCFAH]_i_<[SCFAH]_e_. If internal SCFA are kept low by metabolic activity or transporters, then it would cause a steady influx of SCFAH releasing protons in the more alkaline cytoplasm. Similarly, during fermentation, the increase in [SCFAH]_e_ may slow the passive efflux of newly produced internal SCFAH. In both cases, cells might not be able to balance these fluxes by active proton export, thus causing cytoplasmic acidification at high [SCFAH]_e_. In this scenario, the potency of an SCFA externally added to the medium should scale with its membrane permeability. Contrary to this expectation, ≈ 10× more protonated acetic than lactic acid was required to cause a similar speed decay, Fig. SI15, despite acetic acid having higher membrane permeability through model lipid membranes [36, 46]. This prompted us to look for an alternative explanation.

In both aerobic and anaerobic conditions, assuming fast diffusion of SCFAH across the cell envelope predicts a steady-state internal SCFA concentration 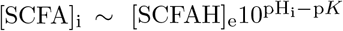. Thus, maintaining pH_i_ ~ 7.5 across the range of [SCFAH]_e_ covered by our experiments would require cells to tolerate large [SCFA]_i_. For example, the threshold lactic acid concentrations below which speeds are not affected, calculated to be [LH] ≈ 0.05 mM (anaerobic) and [LH] ≈ 0.4 mM (aerobic) using the data of Fig. 1(d), would translate respectively to [L]_i_ ~ 275 mM and [L]_i_ ~ 2.20 M, or additional osmotic pressures of respectively 6.7 atm and 53 atm. These pressures far exceed the ~ 0.1 −3 atm previously measured for *E. coli* [47, 48] and should therefore trigger the opening of mechanosensitive channels [49, 50]. Although cells might tolerate internal SCFA and maintain pH_i_ up to the anaerobic threshold simply by releasing internal compounds, there are simply not enough internal solutes that can be exchanged with SCFA to tolerate higher [SCFAH] at constant pH_i_. The failure of pH_i_ homeostasis could, therefore, result from osmotic constraints preventing the maintenance of a transmembrane pH gradient in the presence of SCFAH. This scenario is consistent with the results of Fig. SI15, where the difference between the concentrations of protonated lactic and acetic acids needed to cause a speed decay is consistent with their p*K* difference. Finally, unlike under anaerobic conditions, cells under aerobic conditions can metabolise SCFA, which could delay their intracellular accumulation if the metabolic rate matches or exceeds the rate at which SCFAH cross the membrane. This may explain why [L]_i_ calculated above the aerobic threshold by neglecting degradation and active efflux is unrealistically high.

It is known that growth inhibition by organic acids affects bacterial colonisation of the human colon, a largely anaerobic environment where protonated SCFA are present in high concentrations. For example, human faeces contain ~ 100 mM SCFA [51, 52], and the pH varies between 5.4 and 6.9 in the human colon [52]. 100 mM SCFA with typical p*K* = 4 yields [SCFAH] ≈ 0.3 mM at pH_e_ = 6.5, enough to impair motility and metabolic activity in our experiments. Interestingly, pathogenic enterohaemorrhagic *E. coli* (O157:H7) shows a 1.5-4.3 fold upregulation of motility genes in response to an SCFA mixture mimicking the one found in the small intestine, where pH_e_ is near-neutral and [SCFAH] remains small, and a 23 fold downregulation when exposed to a mixture characteristic of that found in the large intestine [24]. Consequently, it is conceivable that even if motility genes are expressed, motility plays an important role for colonisation only in specific parts of the gastrointestinal tract. Furthermore, *E. coli* relative sensitivity to SCFA challenge may explain why it only amounts to 0.1 − 1% of the bacterial population in the human gut [53, 54], despite growing rapidly in laboratory settings.

## Materials and Methods

### Strains and bacterial culturing

*E. coli* K12 strain AB1157 and its derivatives were used throughout this work. A smooth-swimming mutant (AB1157 Δ*cheY*), a non-motile mutant (AB1157 Δ*fliF*), and a fluorescent strain (AB1157 Δ*flu* expressing eGFP on plasmid pWR21 [55] referred to as AD88) were used as specified. For motility experiments, overnight cultures were inoculated with a single colony and grown for ≈ 16 h in 10 mL LB (37 °C, 200 rpm). The next day, the overnight suspension was diluted 1:100 in tryptone broth (TB: 10 g L^−1^ tryptone, 5 g L^−1^ NaCl), and grown at 30 °C, with shaking at 200 rpm, for 4 h. For pH_i_ measurements, AB1157 strain harbouring the *fliC*^*sticky*^ mutation [56] was transformed with plasmid pkk223-3-pHluorin(M153R) [57] encoding the pH-sensitive fluorescent protein pHluorin. A frozen stock was diluted 1:1000 in LB medium and bacteria were grown at 37 °C with shaking at 220 rpm to OD ≈ 2.0.

### Sample preparation

#### Motility measurements

Cultures grown for motility measurements as described above were washed three times by filtration [31] in 35 mL 1× or 10× Motility Buffer (1× MB: 6.2 mM K_2_HPO_4_, 3.8 mM KH_2_PO_4_, 67 mM NaCl, and 0.1 mM EDTA, pH = 7.0). 10× MB is identical to 1× MB except for the phosphate concentrations: 62 mM K_2_HPO_4_, and 38 mM KH_2_PO_4_. Cells were collected in ~ 1 mL MB after the third filtration (OD ~ 25 − 30). For dense suspension experiments, several cultures were grown and washed simultaneously, and aliquots were merged to produce a single concentrated sample. The concentrated suspension was diluted with motility buffer according to experimental requirements. When cells of strain AD88 were used as tracers for visualising collective motion, AD88 was grown as motile cells, vortexed to make AD88 cells non-motile by shearing their filaments, washed in buffer, and mixed with the dense suspension of motile cells to a final OD of AD88 cells equal to 0.3.

Glucose was added from a 20% (w/v) stock solution to achieve a final glucose concentration of 20 mM, unless otherwise specified. Rectangular glass capillaries (50 × 8 × 0.4mm^3^, Vitrocom) were filled with bacterial suspensions in MB + glucose and immediately sealed with petroleum jelly (Vaseline).

For experiments conducted in the presence of an externally added SCFA, stock solutions were prepared by dissolving the SCFA in MB and were filter-sterilized using a 0.22 µm syringe filter. Appropriate amounts of stock solutions were added to suspensions of washed cells immediately before experiments.

For the experiments of Fig. 3(d), pH_e_ was adjusted by changing the composition of the phosphate buffer following Table SI3. The total phosphate concentration remained fixed at 10 mM.

#### pH_i_ *measurements*

For internal pH_i_ measurements, cells grown as described above were sheared to shorten flagella [34], and washed three times into 1× MB + 20 mM glucose by centrifugation (8000 g, 2 min). Cells were then immobilised with poly-L-lysine attached to the glass coverslip of a custom-made channel (termed tunnel slide) [34, 58], and incubated with 0.5 µm polystyrene beads (Polysciences) for 10 min. The beads attach to the truncated flagellar filaments are used as markers of individual flagellar motors’ rotation. While the cells were prepared for simultaneous flagellar motor speed and pH_i_ measurements, only pH_i_ measurements were possible because the beads often became stuck immediately upon exposure to SCFA-supplemented MB + glucose. However, despite slightly different growth conditions of these two strains, the timing of the pH_i_ and motility drop were consistent.

#### Extraction of fermentation supernatant

Several sterile microcentrifuge tubes were filled completely with dense suspensions (OD = 15) of bacteria in MB + glucose and immediately closed to prevent additional oxygen from diffusing into the suspension. Tubes were stored at room temperature to mimic conditions during motility experiments. At specific times, a single tube was centrifuged (2 min, 13300 rpm), and the supernatant was transferred to a new sterile tube kept on ice. The content of each tube was filter-sterilised (0.22 µm filter) and analysed on the day or stored at −80 °C.

### Data acquisition

#### Motility measurements

Bacteria were imaged at room temperature using a Nikon Ti inverted microscope with a Nikon phase contrast objective (Plan Fluor 10×/0.3) ≈ 100 µm above the bottom of the capillaries. Movies were recorded at 100 fps for 40 s using an Orca Flash 4.0 sCMOS camera.

#### Visualisation of collective motion

A GFP filter set (LEDFITC-A, Semrock) was used to image the fluorescent AD88 tracer cells with the same microscope setup used for differential dynamic microscopy. Movies were recorded at 100 fps for 40 s and the first 500 images of each movie were projected onto a single image to display the motion of the tracers.

#### pH_e_ *measurements*

The pH of all solutions including fermentation supernatants was measured using an InLab Micro electrode (Mettler Toledo), which requires only ≈ 100 µL of sample. The electrode was connected to a pH meter (FiveEasy, Mettler Toledo) and calibrated daily according to the manufacturer’s instructions.

#### Glucose concentration measurements of fermentation supernatants

GAGO20 glucose assay kits (Sigma) were used according to the manufacturer’s instructions to measure glucose concentrations in supernatants. EDTA interferes with the assay and was omitted from the motility buffers used in those experiments.

#### SCFA quantification in fermentation supernatants

NMR was used to identify the compounds present in the fermentation supernatant. Measurements of a filter-sterilized supernatant collected after 30 min of fermentation and of solutions of a single SCFA or of ethanol or glucose prepared for identification were performed on a 800 MHz Bruker UltraStabilised 4 channel instrument with a 5 mm TCI 1H/13C/15N CryoProbe™.

SCFA concentrations were measured by ion chromatography in fermentation supernatants using a Metrohm 883 Basic plus according to the manufacturer’s IC Application Note No. O-37. Calibration standards for lactate, acetate, formate and succinate at concentrations of 0.5 mM, 1 mM, 2 mM, 5 mM and 10 mM were prepared in MB (pH = 7.0) and measured in the same conditions.

#### pH_i_ *measurements*

pH_i_ was measured with ratiometric pHluorin [42, 59]. The sensor was calibrated *in vivo* as previously described [42], using 40 mM potassium benzoate and 40 mM methylamine hydrochloride to set pH_i_ = pH_e_. The ratio of the sensor’s emission intensities at 395 nm and 475 nm excitation was calibrated against the pH of known buffers. Fluorescence imaging was performed with an iXon Ultra 897 EMCCD camera (Andor). A narrow spectrum 395 nm LED (OptoLED, Cairn Research) was used for the excitation at 395 nm and a neutral white LED with ET470/40x filter (Chroma Technology) was used for the 475 nm excitation. Emission was collected at 520 nm using an ET525/40x filter (Chroma Technology). Images were taken every 90 s with 50 ms exposure time.

To obtain pH_i_ measurements both in aerobic and anaerobic conditions, the measurements were performed for 30 min in the tunnel slide, which was sealed immediately after addition of a relevant buffer. For example, measurements in 1× MB + 20 mM glucose were adjusted to the desired pH_e_ with lactic acid by flowing 1× MB + 20 mM glucose + lactic acid into the channel at *t* ≈ 2 min, and immediately sealing the channel after.

### Data analysis and computational methods

#### Motility measurements

DDM post processing and fitting procedures were as described in [31].

#### SCFA quantification in fermentation supernatants

Fermentation products were identified by comparing NMR chemical shifts and ion chromatography retention times recorded on fermentation supernatants with those recorded on standards prepared with a single SCFA or ethanol. The ion chromatography peak areas measured for these standards varied linearly with the known concentrations, and the resulting calibration curves were used to convert peak areas into absolute concentrations for each SCFA. Raw data are given in Tables SI5-7 and calculated concentrations are presented in Table SI1. The relative composition of the SCFA mixture was almost constant for *t* ≥ 5 min: 56.3% lactic acid, 22.6% formic acid, 14.8% acetic acid, and 6.3% succinic acid, where we have first calculated the relative abundances of each SCFA at each time point for the wild type and Δ*fliF* mutant in 1× MB + 20 mM glucose in Table SI1, and then averaged the values obtained for each SCFA at all times *t* ≥ 5 min. Based on these measurements, we prepared SCFA mixtures in 1× MB 20 mM glucose that closely matched those produced by bacteria, Table SI4. We then used the average relative composition of these mixtures, Table SI4, throughout the work.

Ethanol is not detected by ion chromatography and its absolute concentration could not be determined. Given the weak NMR signal of ethanol compared to SCFA, we estimate that its concentration, [Et], does not exceed 15% of the total SCFA concentration. Thus, [Et]*<* 1.4 mM and [Et]*<* 5 mM throughout experiments conducted in 1× MB + 20 mM glucose and 10× MB + 20 mM glucose, respectively. These concentrations do not affect the swimming speed, Fig. SI7, as expected from previous work [58].

The SCFA concentrations reported in Fig. 2(b) for 10× MB at *t* = 90 min and *t* = 120 min were computed from glucose measurements. First, we observed that glucose and SCFA were respectively consumed and produced at constant rates during the first 60 min of fermentation in 10× MB + 20 mM glucose, Fig. SI5. The ratio of these rates provides the amount of SCFA produced per consumed glucose, assumed constant. Then, we calculated the glucose consumption rates in 10× MB + 40 mM for *t* ≥ 60 min from the glucose measurements of Fig. SI5, and used the previously determined glucose-to-SCFA conversion ratio to obtain the corresponding SCFA production rates. Finally, the SCFA concentrations at *t* = 90 min and *t* = 120 min plotted in Fig. 2(b) were calculated using these calculated SCFA production rates for *t* ≥ 60 min, and the SCFA concentration measured at *t* = 60 min in 10× MB + 20 mM glucose.

#### pH_i_ *measurements*

Fluorescence images of pHluorin were analysed as before [42,58]. Otsu thresholding followed by binary erosion and binary dilation was applied to both 475 nm and 395 nm excitation images. The mean intensity was calculated for each cell in both channels. The ratio of these values after background subtraction was finally converted to pH_i_ values using the previously generated calibration curve.

The results of all experiments are presented in Fig. SI13, where individual curves correspond to the average pH_i_ measured on independent batches of cells (typically 20 *< N*_cells_ *<* 40). The average pH_i_ in aerobic conditions is obtained by averaging data from all cells for 10 min *< t <* 20 min at each pH_e_.

Some curves displayed a step-like decrease in pH_i_ at *t* ≈ 30 min, which is reminiscent of the step-like drop in swimming speed upon transition to anaerobic conditions. We interpreted those sudden decays as a sign that the medium turned anaerobic. No drop was observed at pH_e_ = 6.0 & 5.7, which indicates that cells maintain pH_i_ homeostasis in those conditions. Step-like decreases occurred in most but not all curves for 4.7 ≤ pH_e_ ≤ 5.5, confirming the variability in oxygen depletion times. These are due to the variability in the number of cells in the tunnel slide between different slide preparations, where for each estimated cell concentrations (based on the number of cells in a given field of view and known channel height) the timing of the loss of pH_i_ homeostasis was consistent with DDM measurements of speed drops. Finally, no steps were observed in any of the curves recorded at pH_e_ = 4.5 & 4.3, presumably because pH_i_ had already fully decayed in aerobic conditions. The curves that were selected to estimate the anaerobic pH_i_ are displayed in Fig. SI14. For each pH_e_, the anaerobic pH_i_ was determined by averaging the data from those curves at *t >* 28 min.

#### Analysis of Ref. [28] data

The swimming speeds and pH_i_ measured by Minamino *et al*. [28] were manually extracted and pH_i_ was fitted as a function of pH_e_, Fig. SI16. Using the fitted functions (pH_i_ = 0.3483pH_e_ + 4.7205 for MB + 10 mM lactate, pH_i_ = 0.5615pH_e_ +3.1534 for MB + 10 mM lactate + 34 mM acetate), pH_i_ was obtained at the same pH_e_ used in speed measurements, which allowed swimming speeds to be plotted against pH_i_ in Fig. 4(b).

#### Computational methods

The computational work was performed using MATLAB R2020a, R2022a, R2025a. The main steps of the numerical simulations and of the scripts used to compute [SCFAH] and the pH of solutions are described in the main text and SI. When specified, experimental data were interpolated using shape-preserving piecewise cubic interpolation (*interp1* function with *pchip* method).

## ACKNOWLEDGEMENTS

We would like to thank all members of the Poon and Pilizota labs, as well as Terry Hwa, Georg Fritz, Vic Arcus, Sven van Teeffelen and members of their labs for useful discussions. TP and LLN were supported by the Leverhulme Trust grant 594 RPG-2019-187 and HFSP grant RGP013/2024. TP was supported by EPSRC established career fellowship EP/V03264X/1. LLN was supported by a Research Fellowship from the Royal Commission for the Exhibition of 1851 and by Engineering and Physical Sciences Research Council (EPSRC) Centre for Doctoral Training in Soft Matter and Functional Interfaces (Grant EP/L015536/1).

## AUTHOR CONTRIBUTIONS

JSL, LLN, TP, VAM, and WCKP designed research, AD constructed bacterial strains and CD, EK, JSL, LLN, and VAM performed all other research, EK, JSL, LLN, and VAM analyzed the data, LLN, TP, and WCKP wrote the manuscript, and all authors contributed and commented on the manuscript.

## Footnotes

1 Hereafter, *X*_i_ and *X*_e_ denote intracellular and extracellular variables, respectively. We omit the ‘e’ subscript for SCFA concentrations which, unless otherwise stated, are always extracellular.

