## Supplementary Figures, Text and Tables for "Metabolic feedback during bacterial fermentation is a motility brake"

### 1. Acid dissociation constants

The dissociation constants are tabulated in standard conditions ($25\,^{\circ}\mathrm{C}$ and ionic strength $I = 0$) [1], but vary with the ionic strength of the medium:

$$I = \frac{1}{2}\sum_i [x_i] z_i^2, \qquad \textbf{(SI1)}$$

with $[x_i]$ and $z_i$ the concentration and the charge number of ion $i$ in the medium. For instance, the phosphate buffer in MB ($\mathrm{H_2PO_4^-/HPO_4^{2-}}$) has a tabulated $\mathrm{p}K^{\circ} = -\log(K^{\circ}) = 7.2$, but $\mathrm{p}K = 6.9$ at an ionic strength similar to that of MB [2], $I_{\mathrm{MB}} \approx 0.09\,\mathrm{M}$. The $\mathrm{p}K$ of all acid/base couples (A/B) relevant to this study, Table SI2, were adjusted for the ionic strengths of $1\times$ and $10\times$ MB + $20\,\mathrm{mM}$ glucose using the Davies equation[1] [3]:

$$\mathrm{p}K = \mathrm{p}K^{\circ} - C\left(z_{\mathrm{B}}^2 - z_{\mathrm{A}}^2\right)\left(\frac{\sqrt{I}}{1+\sqrt{I}} - 0.3I\right), \qquad \textbf{(SI2)}$$

with $C \approx 0.509$, and $z_{\mathrm{A}}$, $z_{\mathrm{B}}$ the charges of A and B.

### 2. Computing $\mathrm{pH_e}$

We compute $\mathrm{pH_e}$ in the presence of weak acids by solving a system of equations that includes the dissociation equilibrium of each acid, the conservation of matter, the self-ionization of water and charge balance. For a mixture of lactic, acetic, formic and succinic acid dissolved in a phosphate buffer, this gives 14 equations:

[1]Since adding SCFA can further modify the ionic strength of the solution, these constants were updated for each iteration of the scripts used here.

$$\begin{cases} K_{\mathrm{H_3PO_4}}[\mathrm{H_3PO_4}] = a\left(\mathrm{H_3O^+}\right)[\mathrm{H_2PO_4^-}] & \text{(a)} \\ K_{\mathrm{H_2PO_4^-}}[\mathrm{H_2PO_4^-}] = a\left(\mathrm{H_3O^+}\right)[\mathrm{HPO_4^{2-}}] & \text{(b)} \\ K_{\mathrm{LH}}[\mathrm{LH}] = a\left(\mathrm{H_3O^+}\right)[\mathrm{L^-}] & \text{(c)} \\ K_{\mathrm{AH}}[\mathrm{AH}] = a\left(\mathrm{H_3O^+}\right)[\mathrm{A^-}] & \text{(d)} \\ K_{\mathrm{FH}}[\mathrm{FH}] = a\left(\mathrm{H_3O^+}\right)[\mathrm{F^-}] & \text{(e)} \\ K_{\mathrm{SH_2}}[\mathrm{SH_2}] = a\left(\mathrm{H_3O^+}\right)[\mathrm{SH^-}] & \text{(f)} \\ K_{\mathrm{SH^-}}[\mathrm{SH^-}] = a\left(\mathrm{H_3O^+}\right)[\mathrm{S^{2-}}] & \text{(g)} \\ K_{\mathrm{w}} = a\left(\mathrm{H_3O^+}\right)a\left(\mathrm{HO^-}\right) & \text{(h)} \\ [\mathrm{H_3PO_4}] + [\mathrm{H_2PO_4^-}] + [\mathrm{HPO_4^{2-}}] = c_0 & \text{(i)} \\ [\mathrm{LH}] + [\mathrm{L^-}] = [\mathrm{LH}]_0 & \text{(j)} \\ [\mathrm{AH}] + [\mathrm{A^-}] = [\mathrm{AH}]_0 & \text{(k)} \\ [\mathrm{FH}] + [\mathrm{F^-}] = [\mathrm{FH}]_0 & \text{(l)} \\ [\mathrm{SH_2}] + [\mathrm{SH^-}] + [\mathrm{S^{2-}}] = [\mathrm{SH_2}]_0 & \text{(m)} \\ [\mathrm{K^+}]_0 + [\mathrm{H_3O^+}] = [\mathrm{L^-}] + [\mathrm{A^-}] + [\mathrm{F^-}] + [\mathrm{SH^-}] + [\mathrm{HO^-}] & \\ \qquad + [\mathrm{H_2PO_4^-}] + 2\times\left([\mathrm{S^{2-}}] + [\mathrm{HPO_4^{2-}}]\right) & \text{(n)} \end{cases}, \tag{SI3}$$

where $K_{\mathrm{w}} = 10^{-14}$ is the self-ionization constant of water, $[\mathrm{X}]_0$ are the total concentrations of each SCFA and $c_0$ is the total phosphate concentration (*i.e.* the buffer strength). The third dissociation of phosphoric acid, which has a high p$K$, is omitted. The adjusted dissociation constants, obtained from Eq. SI2 and presented in Table SI2, relate the molar concentrations of an acid/base couple to the activity of hydronium ions: $a\left(\mathrm{H_3O^+}\right) = \gamma_{\mathrm{H_3O^+}}[\mathrm{H_3O^+}] = 10^{-\mathrm{pH_e}}$, where $\gamma_{\mathrm{H_3O^+}}$ is the ionic strength-dependent activity coefficient of $\mathrm{H_3O^+}$. $\gamma_{\mathrm{H_3O^+}}$ and $\gamma_{\mathrm{HO^-}}$ are evaluated using the Davies equation for activity coefficients [3]:

$$\log\left(\gamma_{\mathrm{H_3O^+}}\right) = \log\left(\gamma_{\mathrm{HO^-}}\right) = -Cz^2\left(\frac{\sqrt{I}}{1+\sqrt{I}} - 0.3I\right), \tag{SI4}$$

with $C = 0.509$ and $z^2_{\mathrm{H_3O^+}} = 1$. Eq. SI3(n) enforces charge balance. The concentrations of sodium and chloride ions are equal and constant, therefore their contributions to the total charge cancel out and are omitted. $[\mathrm{K^+}]_0$ is the concentration of potassium ions added while mixing $\mathrm{KH_2PO_4}$ and $\mathrm{K_2HPO_4}$ to prepare the phosphate buffer, and remains constant during an experiment. As $\mathrm{pH_e}$ changes, the charges carried by potassium and phosphoric acid ions do not necessarily cancel out, and both need to be included in the charge balance. For each experimental condition, $[\mathrm{K^+}]_0$ is calculated and fed into Eq. SI3(n) using:

$$\begin{cases} [\mathrm{H_3PO_4}]_0 = \dfrac{10^{-2\mathrm{pH_{e,0}}}\, c_0}{K_{\mathrm{H_3PO_4}}K_{\mathrm{H_2PO_4^-}} + 10^{-2\mathrm{pH_{e,0}}} + K_{\mathrm{H_3PO_4}}10^{-\mathrm{pH_{e,0}}}} & \text{(a)} \\ [\mathrm{H_2PO_4^-}]_0 = \dfrac{[\mathrm{H_3PO_4}]_0 K_{\mathrm{H_3PO_4}}}{10^{-\mathrm{pH_{e,0}}}} & \text{(b)} \\ [\mathrm{HPO_4^{2-}}]_0 = \dfrac{[\mathrm{H_2PO_4^-}]_0 K_{\mathrm{H_2PO_4^-}}}{10^{-\mathrm{pH_{e,0}}}} & \text{(c)} \\ [\mathrm{K^+}]_0 = [\mathrm{H_2PO_4^-}]_0 + 2\times[\mathrm{HPO_4^{2-}}]_0 & \text{(d)} \end{cases}, \tag{SI5}$$

where $\mathrm{pH_{e,0}}$ is the known $\mathrm{pH_e}$ of the medium as prepared. Solving Eq. SI3 numerically using the *vpasolve* function in MATLAB R2020a & R2022a returns the multiple solutions of the system, among which only one is physically possible (other solutions include negative concentrations). This unique solution yields the concentrations of protonated and deprotonated SCFA, and $\mathrm{pH_e}$ is calculated using $\mathrm{pH_e} = -\log(a\left(\mathrm{H_3O^+}\right))$.

We validate the computational results by comparing the experimentally measured $\mathrm{pH_e}$ of $1\times$ MB + $20\,\mathrm{mM}$ glucose supplemented with known amounts of an individual SCFA, with the calculated values computed using Eq. SI3. Calculated $\mathrm{pH_e}$ agreed well with experimentally measured values, Fig. SI10(a)-(d). This was also the case for a SCFA mixture prepared to match the composition of the fermentation supernatant, and for the supernatant of a bacterial suspension, Fig. SI10(e). In these control experiments, the best agreement was consistently obtained for a phosphate concentration $c_0 = 11\,\mathrm{mM}$, and this value was used in Fig. SI10. The small discrepancy with the expected phosphate concentration in $1\times$ MB ($c_0 = 10\,\mathrm{mM}$) can be explained by a small loss of water ($\lesssim 10\%$) during autoclaving or pipetting errors.

### 3. Numerics

We summarise below the main steps of the numerics used in Fig. 3 of the main text to predict the temporal evolution of the swimming speed during fermentation:

1. Initialization: The phosphate and potassium concentrations are computed from the known initial $\mathrm{pH_{e,0}}$ of the medium, using Eq. SI5 with $c_0 = [\mathrm{H_3PO_4}]_0 + [\mathrm{H_2PO_4^-}]_0 + [\mathrm{HPO_4^{2-}}]_0 = 10\,\mathrm{mM}$. Dissociation constants, Table SI2, are adjusted using the Davies equation, Eq. SI2.

2. Temporal evolution of $[\mathrm{SCFAH}]$:

   - The SCFA production rate at the previous time step, $r(t-\delta t)$, is computed from $[\mathrm{SCFAH}](t-\delta t)$ using $r([\mathrm{SCFAH}](t)) = r_1 e^{-[\mathrm{SCFAH}](t)/c_1} + r_2 e^{-[\mathrm{SCFAH}](t)/c_2}$, Fig. SI9.
   - The total SCFA concentration is computed using: $[\mathrm{SCFA}](t) = [\mathrm{SCFA}](t-\delta t) + \mathrm{OD} \times \delta t \times r(t-\delta t)$.
   - The concentration of each SCFA is computed from $[\mathrm{SCFA}]$ using the relative composition of the SCFA mixture: 57.1% lactic acid, 21.7% formic acid, 14.6% acetic acid, and 6.6% succinic acid, Table. SI4.
   - $\mathrm{pH_e}$ and $[\mathrm{SCFAH}](t)$ are computed by solving Eq. SI3.
   - All dissociation constants and activity coefficients are updated for the next time step using the Davies equations.

3. $[\mathrm{SCFAH}]$ is converted into normalised speeds using Eq. 5 of the main text.

4. Normalised speeds are converted into absolute swimming speeds, which can now be plotted against time and compared with experimental results.

**Table SI1.** SCFA quantification in fermentation supernatants by ion chromatography

| Medium | $t$ (min) | AB1157 | | | | | AB1157 $\Delta$*fliF* (non-motile) | | | | |
|---|---|---|---|---|---|---|---|---|---|---|---|
| | | A (mM) | L (mM) | F (mM) | S (mM) | Total (mM) | A (mM) | L (mM) | F (mM) | S (mM) | Total (mM) |
| 1× MB + 20 mM glucose | 1 | 0.12 | 0 | 0 | 0.01 | 0.13 | 0.25 | 0 | 0.01 | 0.13 | 0.40 |
| | 5 | 0.36 | 1.33 | 0.59 | 0.08 | 2.36 | 0.46 | 0.88 | 0.49 | 0.20 | 2.03 |
| | 10 | 0.64 | 2.21 | 0.94 | 0.16 | 3.95 | 0.54 | 2.61 | 0.80 | 0.34 | 4.30 |
| | 15 | 0.65 | 3.08 | 1.11 | 0.25 | 5.08 | 0.70 | 3.76 | 1.13 | 0.47 | 6.06 |
| | 20 | 0.73 | 3.53 | 1.37 | 0.32 | 5.94 | 0.96 | 4.11 | 1.44 | 0.53 | 7.05 |
| | 25 | 0.87 | 3.72 | 1.62 | 0.37 | 6.58 | | | | | |
| | 30 | 1.11 | 3.77 | 1.86 | 0.42 | 7.16 | 1.12 | 4.46 | 1.72 | 0.55 | 7.84 |
| | 60 | 1.35 | 4.44 | 2.17 | 0.44 | 8.40 | 1.34 | 4.95 | 1.91 | 0.58 | 8.78 |
| 10× MB + 20 mM glucose | 1 | 0.22 | 0 | 0.03 | 0.09 | 0.33 | 0.32 | 0 | 0.03 | 0.19 | 0.53 |
| | 5 | 0.48 | 1.47 | 0.82 | 0.13 | 2.91 | 0.53 | 1.20 | 0.55 | 0.25 | 2.53 |
| | 10 | 0.61 | 2.64 | 1.28 | 0.23 | 4.75 | 0.86 | 3.65 | 1.47 | 0.35 | 6.32 |
| | 15 | 0.85 | 4.73 | 2.01 | 0.36 | 7.95 | 1.19 | 6.09 | 2.33 | 0.49 | 10.10 |
| | 20 | 1.03 | 6.55 | 2.57 | 0.47 | 10.62 | 1.39 | 7.85 | 2.97 | 0.57 | 12.77 |
| | 30 | 1.47 | 9.74 | 3.74 | 0.65 | 15.60 | 2.22 | 12.02 | 4.61 | 0.86 | 19.71 |
| | 60 | 3.68 | 20.57 | 8.32 | 1.39 | 33.96 | 4.08 | 20.39 | 8.57 | 1.41 | 34.46 |

A: acetic acid. L: lactic acid. F: formic acid. S: succinic acid.
To obtain the $t = 1\,\text{min}$ supernatants, suspensions were collected and centrifuged as quickly as possible after sealing the capillary, *i.e.* $0\,\text{min} < t \lesssim 2\,\text{min}$.

**Table SI2.** Tabulated and adjusted p$K$ values.

| Acid/Base couple | p$K^\circ$ | p$K$ (MB) | p$K$ (10× MB) |
|---|---|---|---|
| $H_3PO_4/H_2PO_4^-$ | 2.16 | 2.05 | 2.02 |
| $H_2PO_4^-/HPO_4^{2-}$ | 7.2 | 6.89 | 6.8 |
| $LH/L^-$ | 3.86 | 3.76 | 3.73 |
| $AH/A^-$ | 4.76 | 4.66 | 4.63 |
| $FH/F^-$ | 3.75 | 3.65 | 3.62 |
| $SH_2/SH^-$ | 4.21 | 4.11 | 4.08 |
| $SH^-/S^{2-}$ | 5.64 | 5.33 | 5.24 |

$LH/L^-$: lactic acid. $AH/A^-$: acetic acid. $FH/F^-$: formic acid. $SH_2/SH^-$ and $SH^-/S^{2-}$: succinic acid.

**Table SI3.** Composition of the phosphate buffers used in Fig. 3(d)

| Target pH | $K_2HPO_4$ (mM) | $KH_2PO_4$ (mM) | Measured pH |
|---|---|---|---|
| 5 | 0 | 10 | 4.93 |
| 5.3 | 0.35 | 9.65 | 5.29 |
| 5.5 | 0.5 | 9.5 | 5.47 |
| 5.7 | 0.75 | 9.25 | 5.67 |
| 6 | 1.32 | 8.68 | 5.98 |
| 6.5 | 3.3 | 6.7 | 6.45 |
| 7 | 6.15 | 3.85 | 6.96 |

**Table SI4.** Mixtures prepared in $1\times$ MB + $20\,\mathrm{mM}$ glucose imitating the composition of fermentation supernatants

| | SCFA | | | | | | | | | |
|---|---|---|---|---|---|---|---|---|---|---|
| $t$ (min) | A (mM) | L (mM) | F (mM) | S (mM) | A (%) | L (%) | F (%) | S (%) | Total SCFA (mM) | E (mM) |
| 0 | 0 | 0 | 0 | 0 | - | - | - | - | 0 | 0 |
| 1 | 0.25 | 0 | 0 | 0.08 | 76 | 0 | 0 | 24 | 0.33 | 0 |
| 5 | 0.40 | 1.10 | 0.50 | 0.15 | 19 | 51 | 23 | 7 | 2.15 | 0.20 |
| 10 | 0.55 | 2.40 | 0.85 | 0.25 | 14 | 59 | 21 | 6 | 4.05 | 0.50 |
| 15 | 0.70 | 3.40 | 1.10 | 0.35 | 13 | 61 | 20 | 6 | 5.55 | 0.60 |
| 20 | 0.90 | 3.80 | 1.40 | 0.45 | 14 | 58 | 21 | 7 | 6.55 | 0.80 |
| 30 | 1.10 | 4.20 | 1.75 | 0.50 | 15 | 56 | 23 | 7 | 7.55 | 0.90 |
| Average ($t \geq 5\,\mathrm{min}$): | | | | | 14.6 | 57.1 | 21.7 | 6.6 | | |

A: acetic acid. L: lactic acid. F: formic acid. S: succinic acid. E: ethanol.

**Table SI5.** Ion chromatography calibration raw data, showing retention times and areas of peaks recorded for solutions containing a known concentration of a single SCFA in motility buffer. Seven different peaks were identified and attributed either to the motility buffer or to an individual SCFA.

| | | Peak 1 (buffer) | Peak 2 (buffer) | Peak 3 (succinate) | Peak 4 (lactate) | Peak 5 (formate) | Peak 6 (acetate) | Peak 7 (buffer) |
|---|---|---|---|---|---|---|---|---|
| | | Retention time: 10.41 min | Retention time: 11.5 min | Retention time: 15.5 min | Retention time: 17 min | Retention time: 19 min | Retention time: 20 min | Retention time: 23 min |
| Compound | Concentration (mM) | Area (µS.cm$^{-1}$.min) | Area (µS.cm$^{-1}$.min) | Area (µS.cm$^{-1}$.min) | Area (µS.cm$^{-1}$.min) | Area (µS.cm$^{-1}$.min) | Area (µS.cm$^{-1}$.min) | Area (µS.cm$^{-1}$.min) |
| MB | | 50.867 | 5.627 | 0.005 | 0.000 | 0.000 | 0.000 | 0.142 |
| Lactate | 0.5 | 50.589 | 5.126 | 0.004 | 0.159 | 0.004 | 0.000 | 0.149 |
| Lactate | 1 | 50.658 | 5.158 | 0.000 | 0.334 | 0.000 | 0.000 | 0.139 |
| Lactate | 2 | 50.622 | 5.195 | 0.000 | 0.675 | 0.000 | 0.000 | 0.075 |
| Lactate | 5 | 50.641 | 5.257 | 0.023 | 1.701 | 0.007 | 0.000 | 0.144 |
| Lactate | 10 | 50.814 | 5.257 | 0.000 | 3.399 | 0.009 | 0.000 | 0.133 |
| Acetate | 0.5 | 50.239 | 5.221 | 0.000 | 0.000 | 0.000 | 0.229 | 0.141 |
| Acetate | 1 | 50.401 | 5.396 | 0.000 | 0.000 | 0.000 | 0.500 | 0.147 |
| Acetate | 2 | 50.824 | 5.356 | 0.000 | 0.000 | 0.000 | 0.938 | 0.137 |
| Acetate | 5 | 50.413 | 5.338 | 0.000 | 0.000 | 0.000 | 2.426 | 0.141 |
| Acetate | 10 | 50.769 | 5.591 | 0.003 | 0.004 | 0.005 | 4.872 | 0.131 |
| Formate | 0.5 | 50.658 | 5.477 | 0.000 | 0.000 | 0.278 | 0.000 | 0.144 |
| Formate | 1 | 50.323 | 5.437 | 0.000 | 0.000 | 0.552 | 0.000 | 0.144 |
| Formate | 2 | 50.707 | 5.752 | 0.000 | 0.000 | 1.111 | 0.000 | 0.154 |
| Formate | 5 | 50.380 | 5.574 | 0.000 | 0.000 | 2.813 | 0.000 | 0.143 |
| Formate | 10 | 50.413 | 5.580 | 0.000 | 0.000 | 5.577 | 0.000 | 0.139 |
| Succinate | 0.5 | 50.611 | 5.832 | 0.509 | 0.000 | 0.000 | 0.000 | 0.140 |
| Succinate | 1 | 50.606 | 5.759 | 1.063 | 0.000 | 0.000 | 0.000 | 0.141 |
| Succinate | 2 | 50.557 | 5.753 | 2.157 | 0.000 | 0.000 | 0.000 | 0.148 |
| Succinate | 5 | 50.677 | 5.923 | 5.554 | 0.000 | 0.000 | 0.000 | 0.141 |
| Succinate | 10 | 50.328 | 5.764 | 10.726 | 0.000 | 0.000 | 0.000 | 0.138 |

**Table SI6.** Ion chromatography raw data for supernatants harvested from dense suspensions (OD=15) in 1×MB + 20 mM glucose at various time points (1-60 min). The different peaks recorded for the various supernatants matched those of Table. SI5. Supernatants were diluted 2× before the measurement. Peak areas were converted to SCFA concentrations in Table SI1 using this dilution factor and the calibration results of Table. SI5.

| | | | Peak 1 (buffer) | Peak 2 (buffer) | Peak 3 (succinate) | Peak 4 (lactate) | Peak 5 (formate) | Peak 6 (acetate) | Peak 7 (buffer) |
|---|---|---|---|---|---|---|---|---|---|
| | | | Retention time: 10.4 min | Retention time: 11.5 min | Retention time: 15.6 min | Retention time: 17.1 min | Retention time: 19 min | Retention time: 20.1 min | Retention time: 23.1 min |
| Strain | Fermentation time (min) | Dilution factor | Area (µS.cm$^{-1}$.min) | Area (µS.cm$^{-1}$.min) | Area (µS.cm$^{-1}$.min) | Area (µS.cm$^{-1}$.min) | Area (µS.cm$^{-1}$.min) | Area (µS.cm$^{-1}$.min) | Area (µS.cm$^{-1}$.min) |
| AB1157 | 1 | 2 | 48.745 | 5.017 | 0.004 | 0 | 0 | 0.029 | 0.141 |
| AB1157 | 5 | 2 | 48.198 | 5.286 | 0.045 | 0.226 | 0.164 | 0.088 | 0.136 |
| AB1157 | 10 | 2 | 48.449 | 5.318 | 0.089 | 0.375 | 0.262 | 0.156 | 0.143 |
| AB1157 | 15 | 2 | 48.759 | 5.261 | 0.133 | 0.523 | 0.311 | 0.157 | 0.137 |
| AB1157 | 20 | 2 | 48.217 | 5.477 | 0.174 | 0.599 | 0.382 | 0.177 | 0.137 |
| AB1157 | 25 | 2 | 48.368 | 5.169 | 0.202 | 0.631 | 0.452 | 0.212 | 0.137 |
| AB1157 | 30 | 2 | 48.493 | 5.501 | 0.228 | 0.641 | 0.519 | 0.270 | 0.138 |
| AB1157 | 60 | 2 | 48.414 | 5.409 | 0.238 | 0.754 | 0.606 | 0.328 | 0.136 |
| AB1157 Δ*fliF* | 1 | 2 | 50.777 | 5.250 | 0.071 | 0 | 0.004 | 0.061 | 0.140 |
| AB1157 Δ*fliF* | 5 | 2 | 50.439 | 5.339 | 0.107 | 0.150 | 0.138 | 0.111 | 0.136 |
| AB1157 Δ*fliF* | 10 | 2 | 47.435 | 4.829 | 0.186 | 0.444 | 0.224 | 0.132 | 0.127 |
| AB1157 Δ*fliF* | 15 | 2 | 50.885 | 5.242 | 0.256 | 0.638 | 0.315 | 0.170 | 0.138 |
| AB1157 Δ*fliF* | 20 | 2 | 50.605 | 5.368 | 0.287 | 0.699 | 0.403 | 0.233 | 0.138 |
| AB1157 Δ*fliF* | 30 | 2 | 50.813 | 5.333 | 0.295 | 0.757 | 0.481 | 0.272 | 0.138 |
| AB1157 Δ*fliF* | 60 | 2 | 50.541 | 5.329 | 0.313 | 0.841 | 0.532 | 0.327 | 0.137 |

**Table SI7.** Ion chromatography raw data for supernatants harvested from dense suspensions (OD=15) in $10\times$MB + 20 mM glucose at various time points (1-60 min). The different peaks recorded for the various supernatants matched those of Table. SI5. Supernatants were diluted $2\times$ before the measurement. Peak areas were converted to SCFA concentrations in Table SI1 using this dilution factor and the calibration results of Table. SI5.

| | | | Peak 1 (buffer) | Peak 2 (buffer) | Peak 3 (succinate) | Peak 4 (lactate) | Peak 5 (formate) | Peak 6 (acetate) | Peak 7 (buffer) |
|---|---|---|---|---|---|---|---|---|---|
| | | | Retention time: 10.4 min | Retention time: 12.2 min | Retention time: 15.8 min | Retention time: 17.1 min | Retention time: 19 min | Retention time: 20.2 min | Retention time: 23.2 min |
| Strain | Fermentation time (min) | Dilution factor | Area ($\mu$S.cm$^{-1}$.min) | Area ($\mu$S.cm$^{-1}$.min) | Area ($\mu$S.cm$^{-1}$.min) | Area ($\mu$S.cm$^{-1}$.min) | Area ($\mu$S.cm$^{-1}$.min) | Area ($\mu$S.cm$^{-1}$.min) | Area ($\mu$S.cm$^{-1}$.min) |
| AB1157 | 1 | 2 | 48.605 | 55.001 | 0.049 | 0 | 0.007 | 0.053 | 0.136 |
| AB1157 | 5 | 2 | 48.852 | 56.476 | 0.072 | 0.250 | 0.229 | 0.117 | 0.128 |
| AB1157 | 10 | 2 | 48.787 | 55.930 | 0.124 | 0.448 | 0.357 | 0.148 | 0.131 |
| AB1157 | 15 | 2 | 48.740 | 56.309 | 0.196 | 0.804 | 0.560 | 0.207 | 0.130 |
| AB1157 | 20 | 2 | 48.887 | 55.049 | 0.254 | 1.113 | 0.717 | 0.251 | 0.129 |
| AB1157 | 30 | 2 | 48.715 | 54.221 | 0.351 | 1.654 | 1.045 | 0.358 | 0.128 |
| AB1157 | 60 | 2 | 48.730 | 55.848 | 0.748 | 3.494 | 2.324 | 0.896 | 0.126 |
| AB1157 Δ*fliF* | 1 | 2 | 49.350 | 51.450 | 0.104 | 0 | 0.007 | 0.077 | 0.133 |
| AB1157 Δ*fliF* | 5 | 2 | 49.402 | 50.837 | 0.135 | 0.204 | 0.153 | 0.129 | 0.131 |
| AB1157 Δ*fliF* | 10 | 2 | 49.022 | 50.170 | 0.190 | 0.620 | 0.410 | 0.208 | 0.132 |
| AB1157 Δ*fliF* | 15 | 2 | 49.575 | 50.865 | 0.265 | 1.035 | 0.650 | 0.289 | 0.132 |
| AB1157 Δ*fliF* | 20 | 2 | 49.505 | 50.447 | 0.308 | 1.333 | 0.830 | 0.337 | 0.130 |
| AB1157 Δ*fliF* | 30 | 2 | 49.445 | 50.922 | 0.465 | 2.043 | 1.287 | 0.539 | 0.130 |
| AB1157 Δ*fliF* | 60 | 2 | 49.520 | 51.384 | 0.763 | 3.465 | 2.392 | 0.993 | 0.130 |

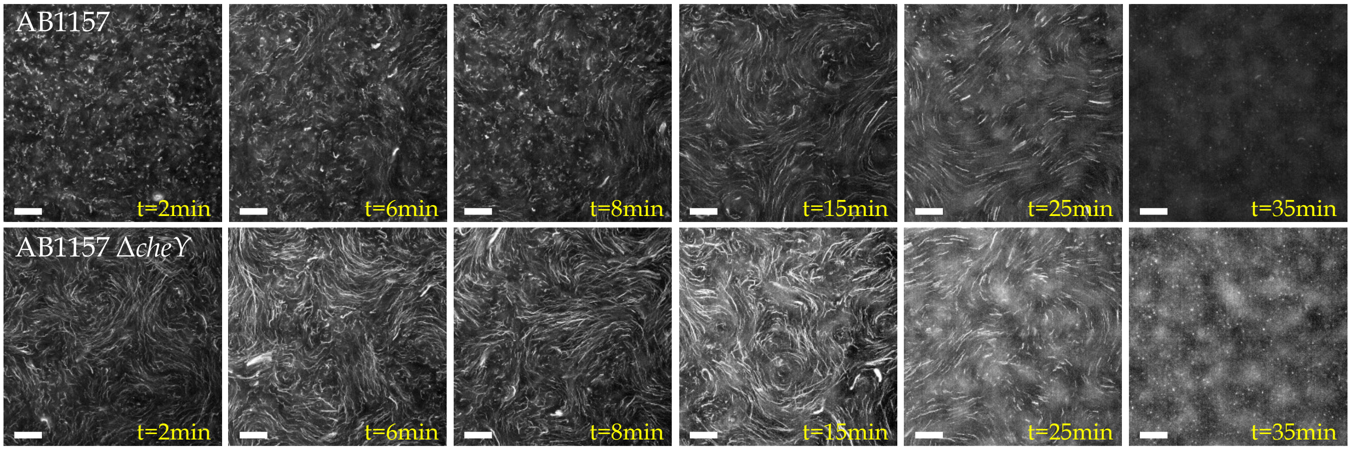


**Fig. SI1.** Collective motion in dense suspensions of *E. coli* (OD=15). Still images are produced by superimposing the first 500 frames of each movie, in which fluorescent non-motile cells of strain AD88 are used as flow tracers (*Methods*). AD88 cells in a suspension of wild type AB1157 *E. coli* (top row), and the smooth-swimming AB1157 $\Delta$*cheY* (bottom row) are shown. In the absence of collective motion, tracers display diffusive motion (for example, $t = 2\,\mathrm{min}$ in the wild type suspension). Collective motion is evident from the ballistic trajectories displayed by the non-motile tracers from $t = 8\,\mathrm{min}$ in the wild type suspension and from the start of the experiment in the suspension of smooth-swimming *E. coli*. Collective motion has previously been shown to enhance the swimming speed measured in dense suspensions of *E. coli* by DDM, especially for cell body volume fraction $\phi \gtrsim 0.05$ (although in tighter confinement) [4]. Considering a cell concentration $c \sim 2.5 \times 10^{10}$ cells/mL and a cell volume $V_{\mathrm{cell}} \approx 2.3\,\mathrm{fL}$ [5], we estimate $\phi \approx 0.06$ at $\mathrm{OD} = 15$ in our experiments and attribute the increased speed recorded at OD=15 to the emergence of collective motion. Scale bars: $50\,\mathrm{\mu m}$.

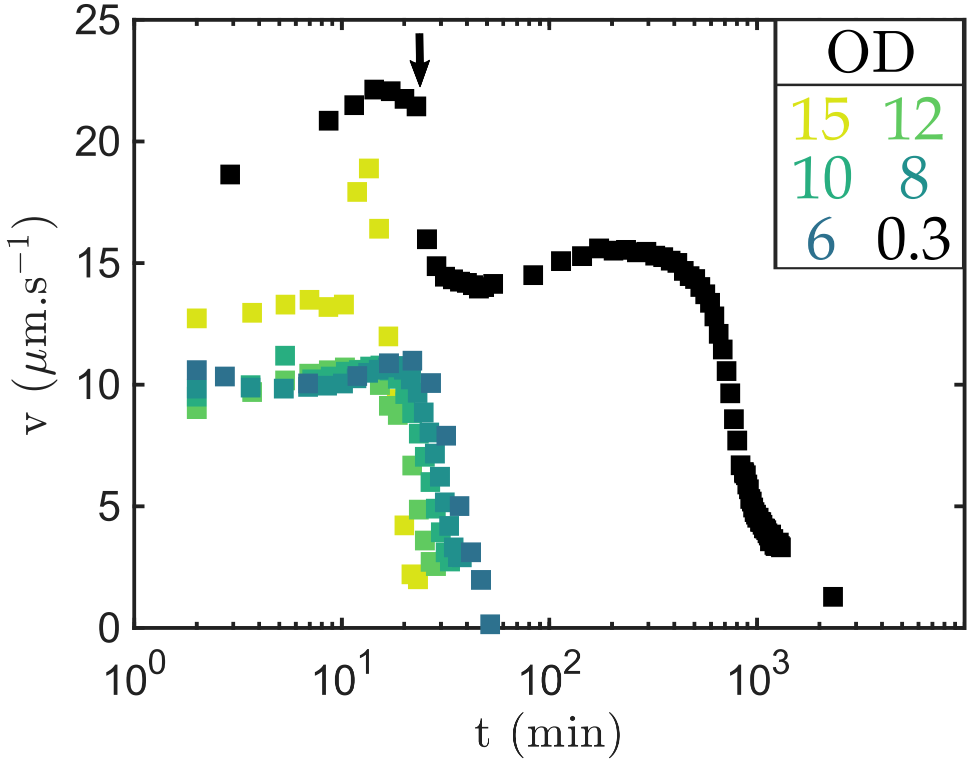


**Fig. SI2.** Swimming speeds obtained as a function of time for the conditions presented in Fig. 1(b) of the main text, with the addition of the results for a dilute suspension (OD=0.3, black). The first speed decay seen at OD=0.3 ($t \approx 20\,\mathrm{min}$, black arrow) corresponds to the transition from aerobic to anaerobic conditions. This transition occurs at $t < 2\,\mathrm{min}$ for the denser suspensions and is thus not visible.

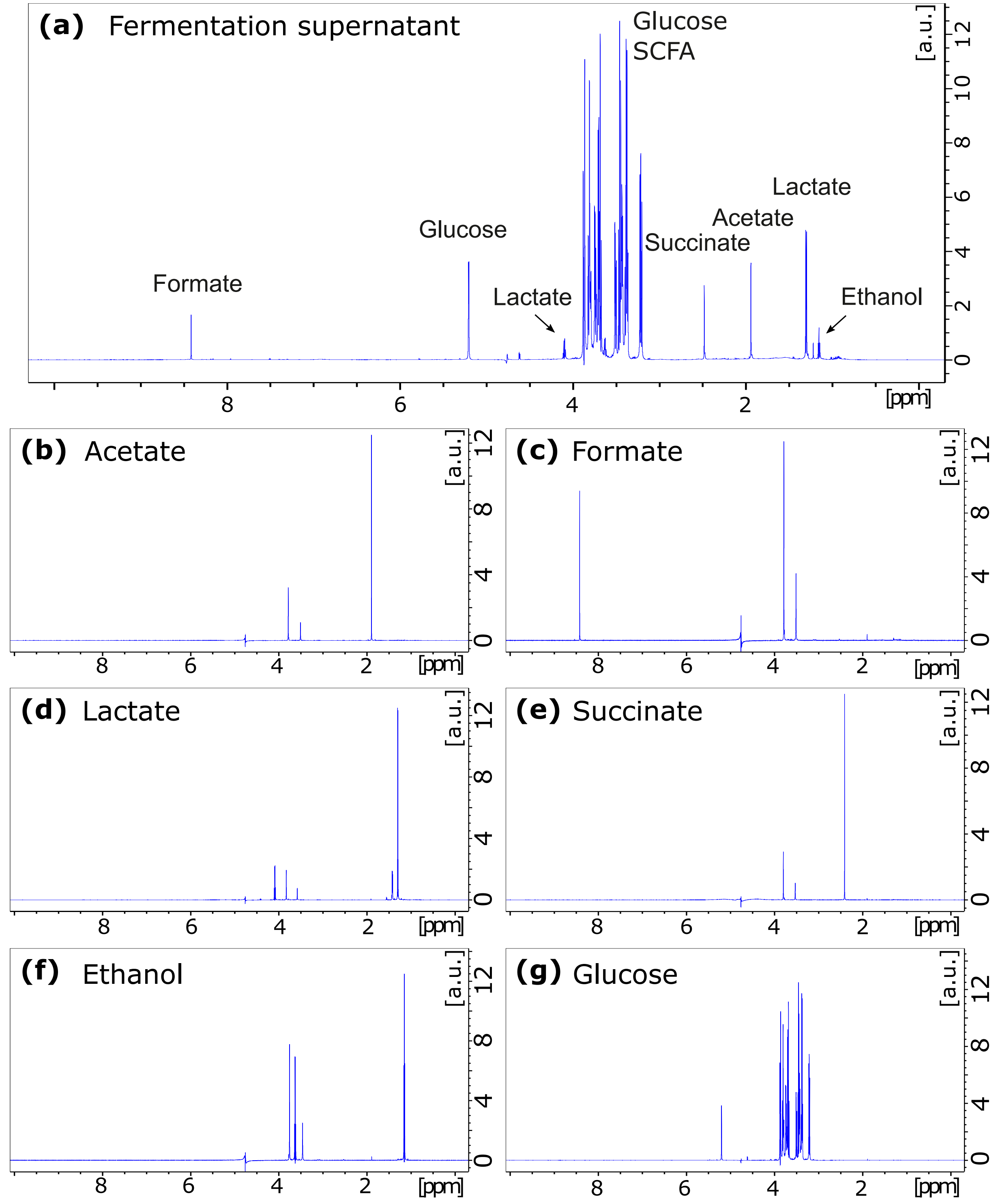


**Fig. SI3.** **(a)** Proton NMR spectrum recorded on a dense suspension supernatant after $30\,\mathrm{min}$ of fermentation. The peaks are attributed to specific species by comparison with reference spectra recorded for individual compounds and presented in **(b)-(g)**.

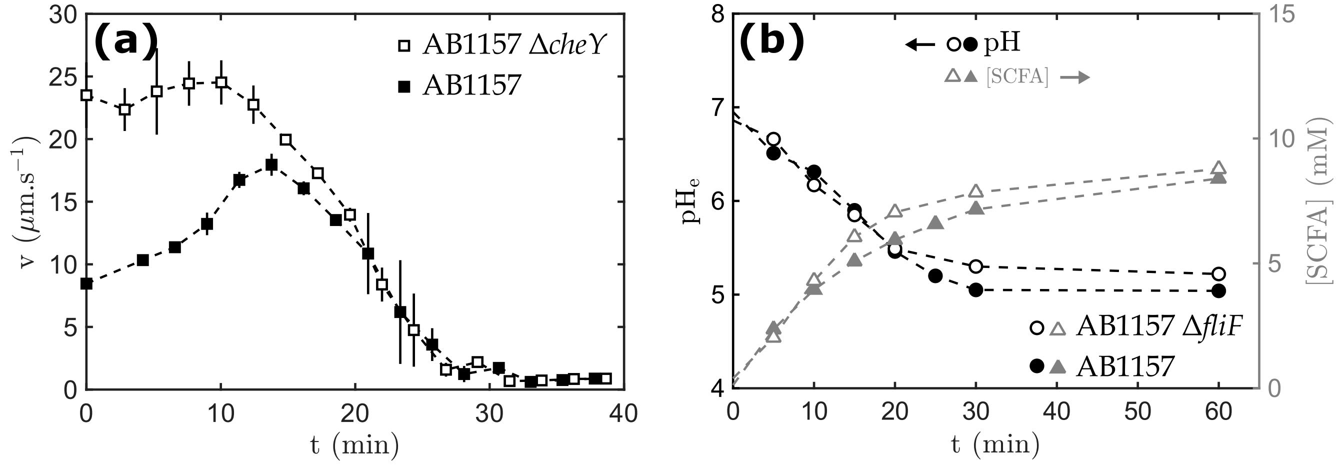


**Fig. SI4.** *E. coli* strains with mutations (open markers) in motility genes display physiological perturbations identical to those exhibited by the wild type (filled markers). **(a)** Dense suspensions (OD=15) of wild type (filled squares) and smooth-swimming ($\Delta$*cheY*, open squares) AB1157 display similar motility decay after $10\text{-}25\,\mathrm{min}$. The $\Delta$*cheY* strain shows an immediate and more pronounced speed enhancement compared to the wild type, for which collective motion begins at $t \approx 10\,\mathrm{min}$ (see also Fig. SI1). Error bars are standard error of the mean of two independent experiments. **(b)** Dense suspensions (OD=15) of the wild type (filled markers) and non-motile ($\Delta$*fliF*, open markers) strains. Both strains secrete similar amounts of SCFA during fermentation (gray triangles), causing an identical decrease in the $\mathrm{pH_e}$ of the medium (black circles).

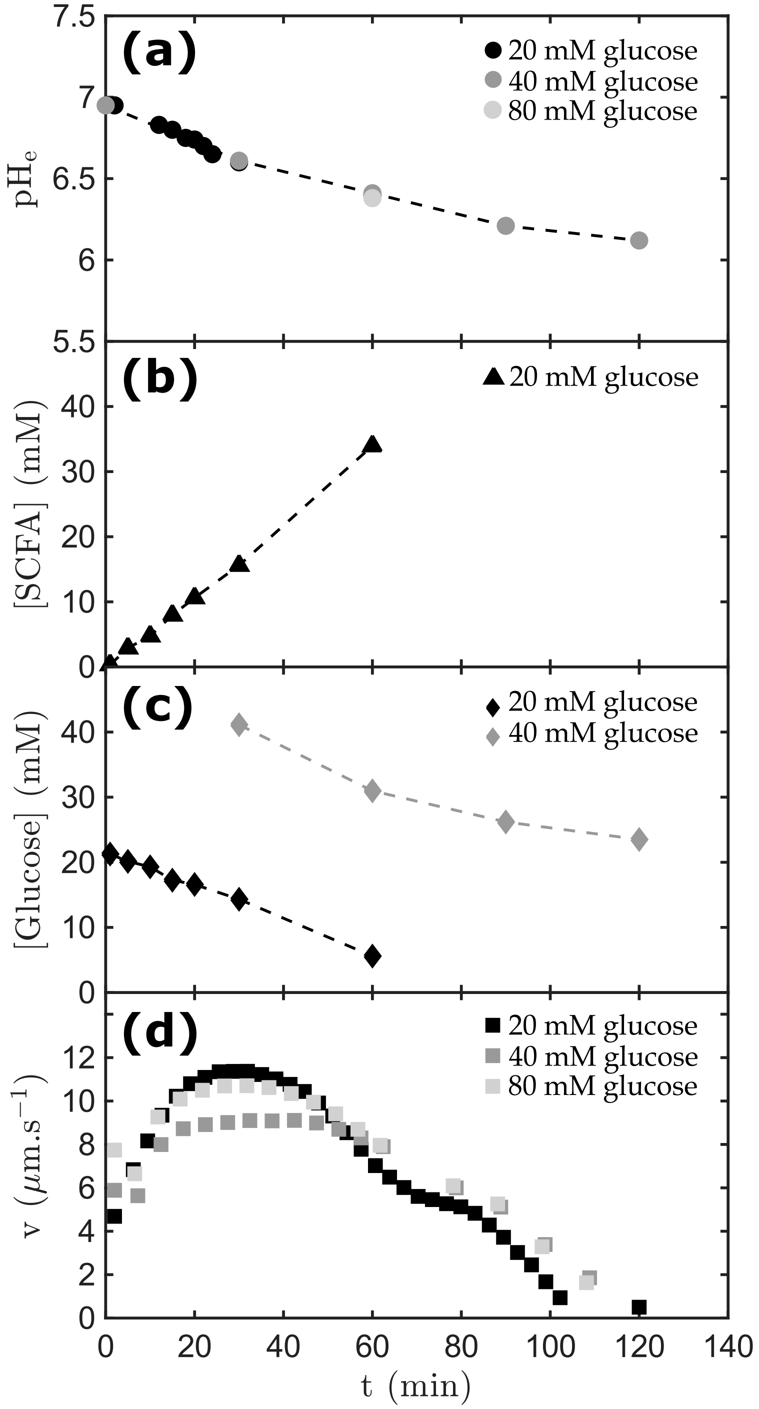


**Fig. SI5.** Metabolic activity and speed decay for dense suspensions (OD=15) in $10\times$ MB supplemented with $20\,\mathrm{mM}$ (black), $40\,\mathrm{mM}$ (dark grey), or $80\,\mathrm{mM}$ (light grey) glucose. **(a)** $\mathrm{pH_e}$ (circles) as a function of time. **(b)** SCFA concentration (triangles) in the medium as a function of time. **(c)** Glucose concentration (diamonds) in the medium as a function of time. **(b)** Swimming speed (squares) as a function of time. Speed variations are not affected by the initial glucose concentration, showing that the speed decay is not dictated by the amount of glucose. Data from panels (b) and (c) were used to estimate the SCFA concentrations presented in Fig. 2(b) in the main text at $t = 90\,\mathrm{min}$ $t = 120\,\mathrm{min}$ (see *Methods*).

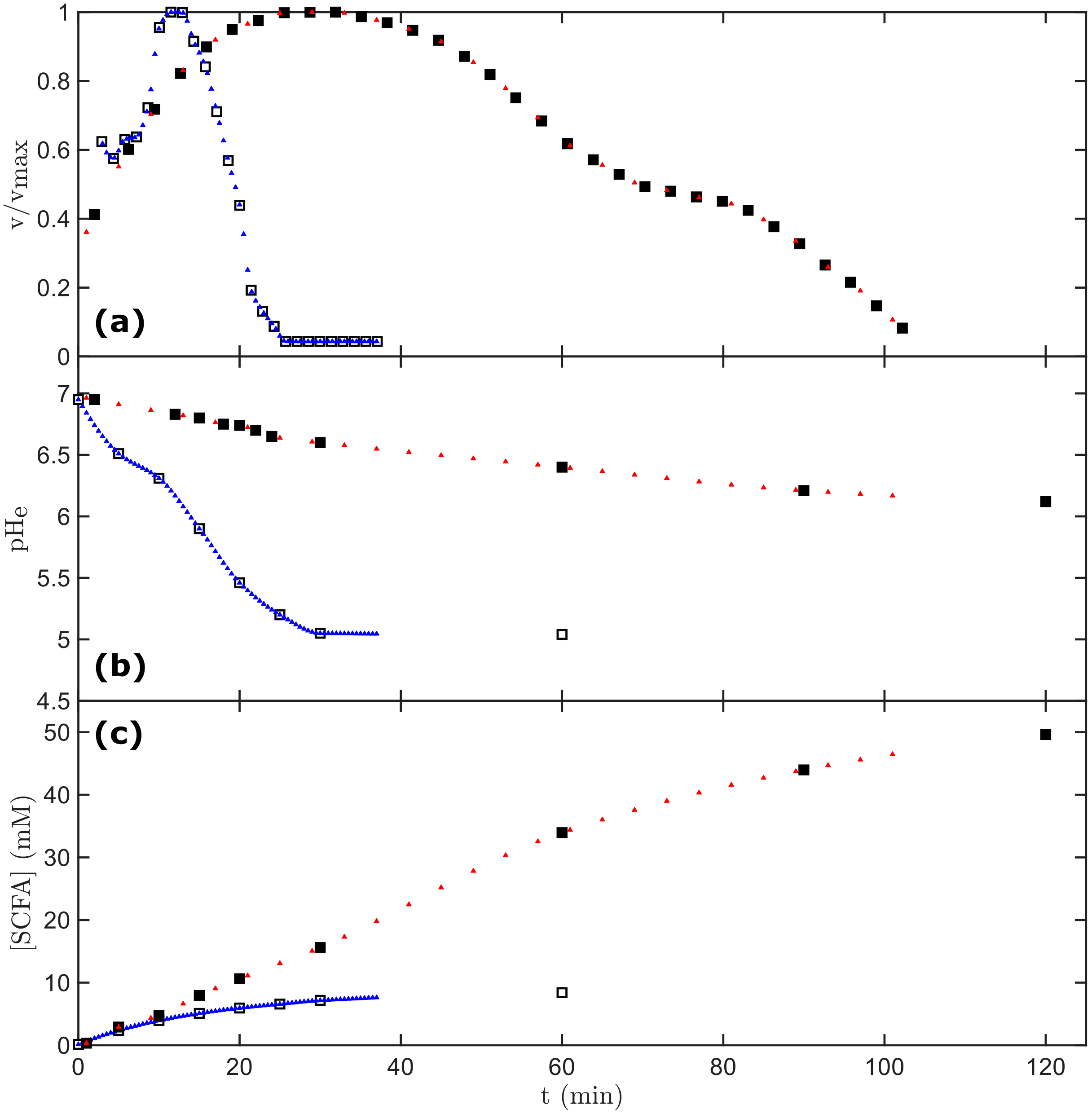


**Fig. SI6.** Interpolation of experimentally measured swimming speeds (normalised to the $v_{\mathrm{max}}$) in **(a)**, $\mathrm{pH_e}$ in **(b)**, and [SCFA] in **(c)**, all depicted with squares. We used shape-preserving piecewise cubic interpolation (*Methods*) to obtain the three variables at the same time points, where the interpolated values are given as triangles. $0.5\,\mathrm{min}$ and $4\,\mathrm{min}$ time steps were used to interpolate the $1\times$ MB + $20\,\mathrm{mM}$ glucose (open squares and blue triangles), and $10\times$ MB + $20\,\mathrm{mM}$ glucose (filled squares and red triangles) results, respectively.

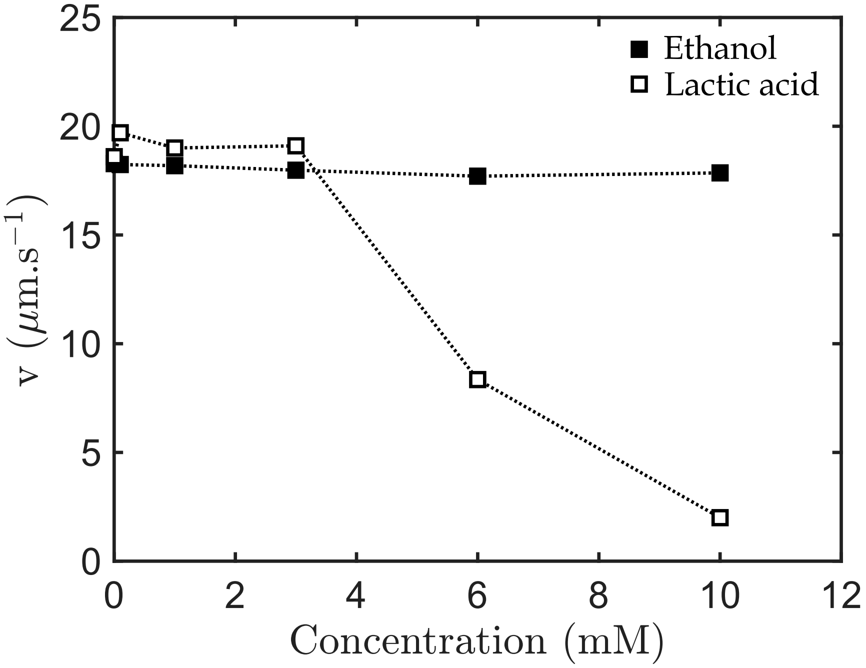


**Fig. SI7.** Swimming speeds measured in dilute suspensions (OD=0.3) after the transition to anaerobic conditions ($t = 60\,\mathrm{min}$) in $1\times$ MB + $20\,\mathrm{mM}$ glucose, and in the presence of the given ethanol (filled squares) or lactic acid (open squares) concentration added to the medium. Whereas lactic acid supplementation causes the speed to decay, ethanol has no effect in the range of concentrations tested, which exceeds the range of concentrations produced by the bacteria in our experiments.

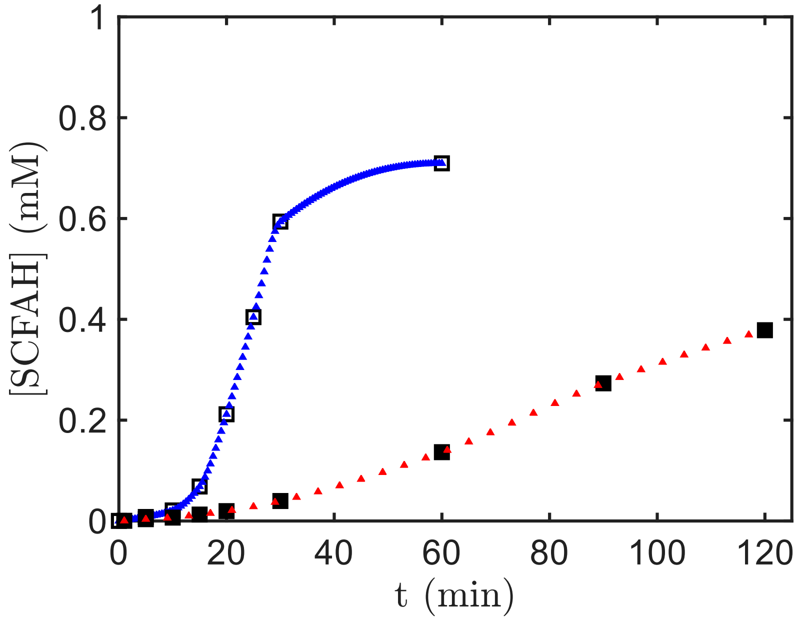


**Fig. SI8.** Concentration of uncharged organic acids as a function of time in $1\times$ MB + $20\,\mathrm{mM}$ glucose (open squares and blue triangles) and $10\times$ MB + $20\,\mathrm{mM}$ glucose (filled squares and red triangles). Squares depict the total [SCFAH] calculated from Eq. 1 and Eq. 2 of the main text, using the experimentally measured [SCFA], $\mathrm{pH_e}$ of Fig. 2(b) and the average relative composition of the SCFA mixture as inputs. Triangles depict the interpolated values, using the same interpolated time vector as in Fig. SI6.

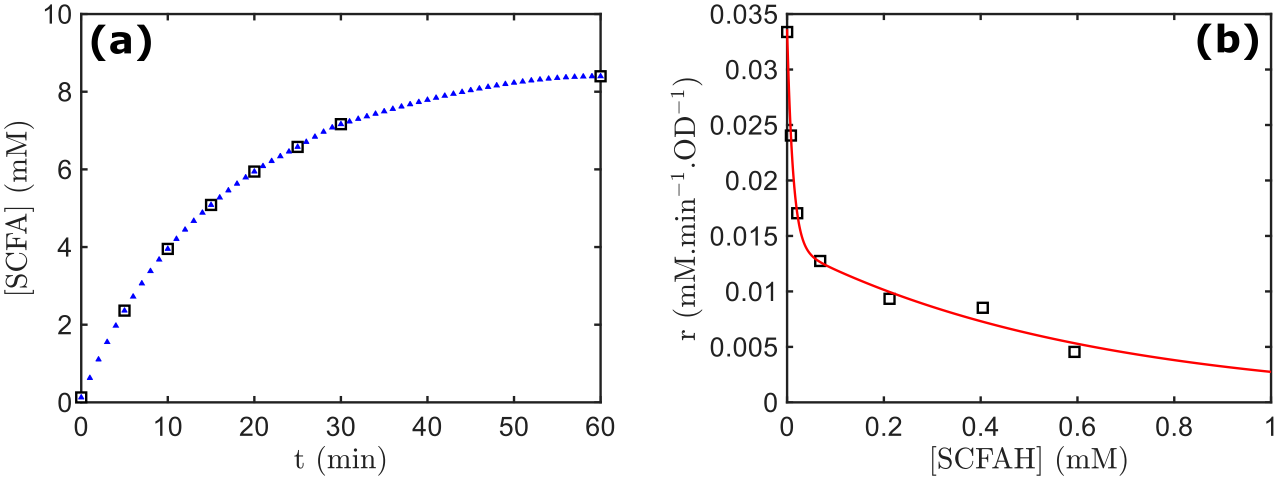


**Fig. SI9.** Empirical relationship for the SCFA production rate in MB as a function of [SCFAH]. **(a)** Squares: [SCFA] plotted as a function of time for a dense (OD=15) suspension in $1\times$ MB + $20\,\mathrm{mM}$ glucose (data from Fig. 2(b) of the main text). Triangles: Interpolated values obtained using shape-preserving piecewise cubic interpolation and $1\,\mathrm{min}$ time steps. **(b)** Squares: SCFA production rate plotted as a function of [SCFAH]. Rates are calculated as the [SCFA] increment between an experimental data point and the next interpolated point in panel (a), divided by the time step ($\delta t = 1\,\mathrm{min}$)) and by the cell density (OD=15). Each rate is plotted as a function of the corresponding [SCFAH], already calculated in Fig. SI8. Red line: fitted function $r([\mathrm{SCFAH}]) = r_1 e^{-[\mathrm{SCFAH}]/c_1} + r_2 e^{-[\mathrm{SCFAH}]/c_2}$, with $r_1 = 0.020\,\mathrm{mM\,min^{-1}\,OD^{-1}}$, $r_2 = 0.014\,\mathrm{mM\,min^{-1}\,OD^{-1}}$, $c_1 = 0.012\,\mathrm{mM}$ and $c_2 = 0.61\,\mathrm{mM}$.

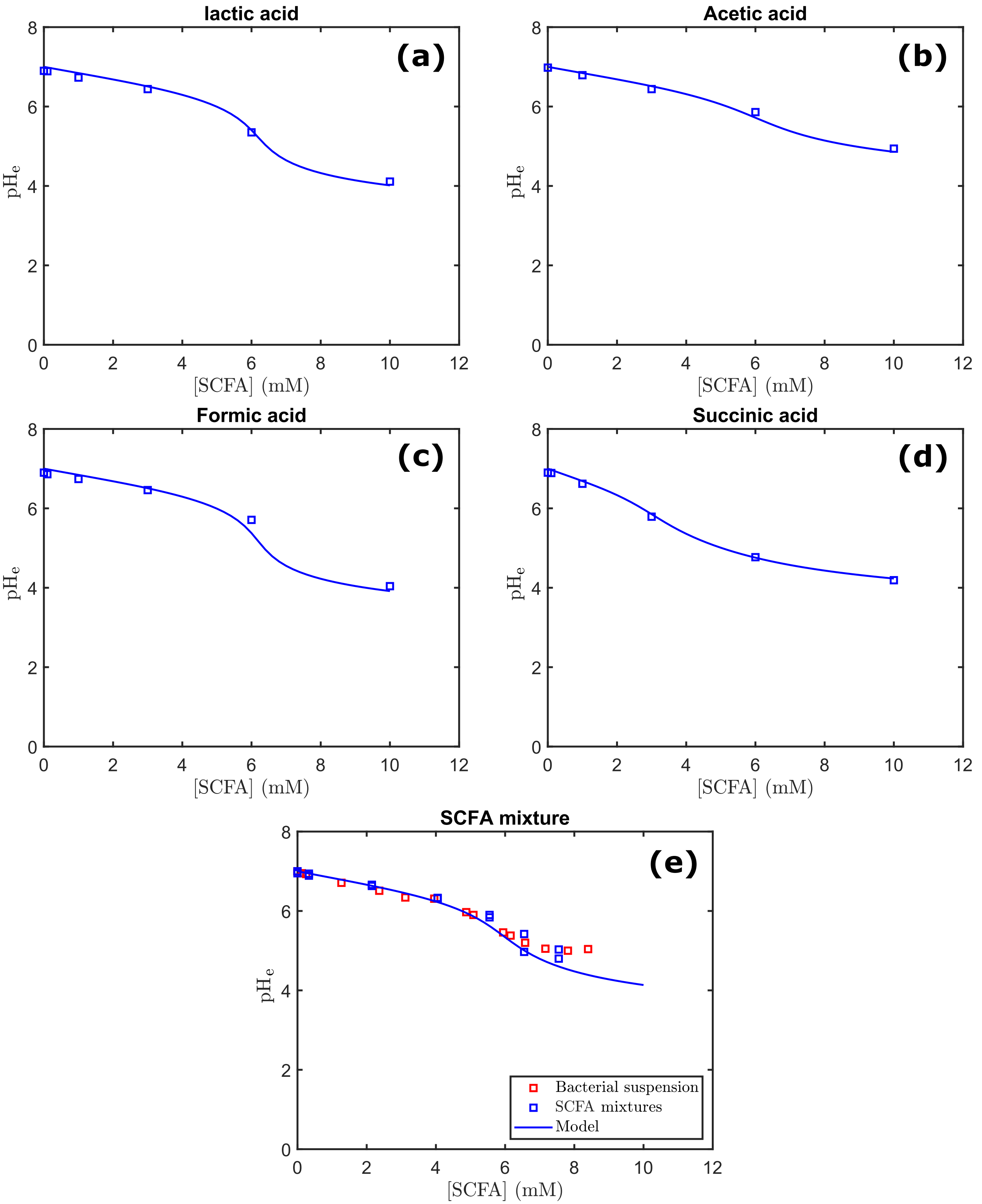


**Fig. SI10.** Evolution of the $\mathrm{pH_e}$ as a function of the total SCFA concentration. Blue squares: experimental results obtained for $1\times$ MB + $20\,\mathrm{mM}$ glucose prepared to known SCFA concentrations (see also Table SI4). Red squares: experimental results obtained for the supernatants of dense bacterial suspensions (OD=15) in $1\times$ MB + $20\,\mathrm{mM}$ glucose, for which [SCFA] was measured by ion chromatography. Blue lines: $\mathrm{pH_e}$ values calculated by solving Eq. SI3, with $\mathrm{pH_{e,0}} = 7.0$ and $c_0 = 11\,\mathrm{mM}$. **(a)** Lactic acid. **(b)** Acetic acid. **(c)** Formic acid. **(d)** Succinic acid. **(e)** SCFA mixture containing all SCFA, obtained either by harvesting the supernatant of a bacterial suspension (red) or by mixing SCFA in MB according to the proportions measured by ion chromatography (blue, two independent repeats), Table SI4.

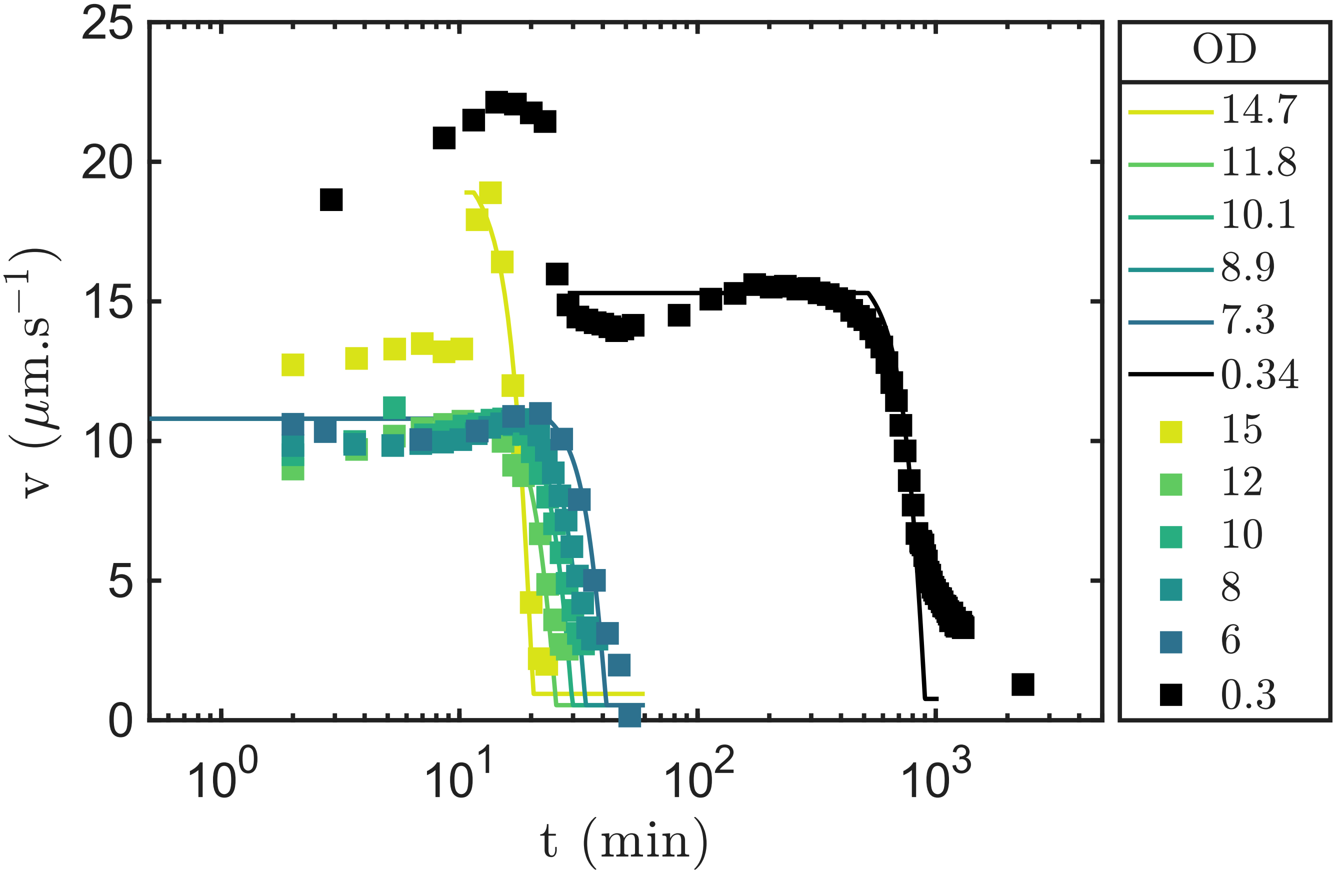


**Fig. SI11.** Experimental (squares) and simulation (continuous lines) results obtained for bacterial suspensions with a range of ODs. Identical to Fig. 3(c) of the main text, with the addition of the experimental data obtained at OD=0.3 and calculated speeds. OD values are indicated in the legend on the right.

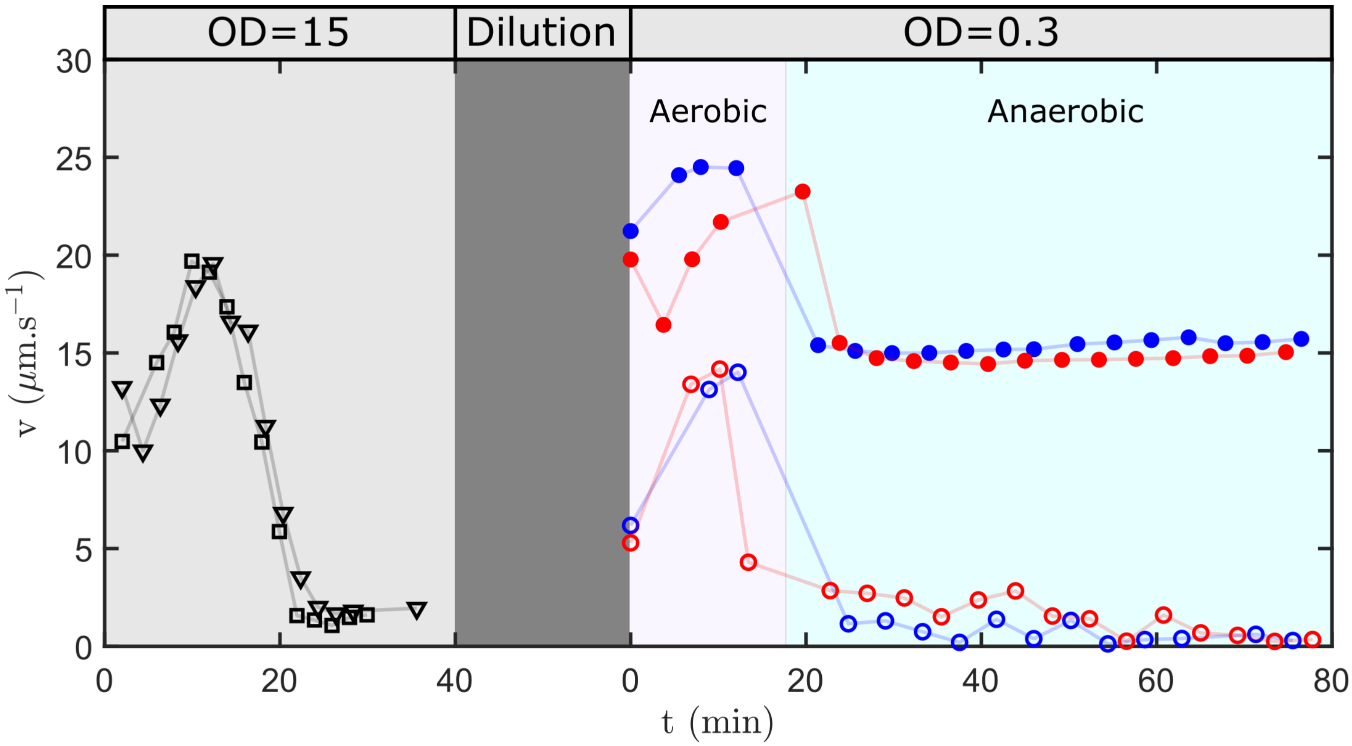


**Fig. SI12.** Effect of a fermentation supernatant collected from dense (OD=15), sealed suspensions on swimming speeds of bacteria in different conditions. Red markers depict bacteria diluted to OD=0.3 in AB30 (empty circles) or $1\times$ MB + $20\,\mathrm{mM}$ glucose (filled circles), where AB30 is the supernatant obtained after $\approx 30\,\mathrm{min}$ of fermentation in $1\times$ MB + $20\,\mathrm{mM}$ glucose at OD=15. Blue markers depict cells harvested after $\approx 50\,\mathrm{min}$ of fermentation in $1\times$ MB + $20\,\mathrm{mM}$ glucose at OD=15, and then diluted to OD=0.3 in AB30 (empty circles) or $1\times$ MB + $20\,\mathrm{mM}$ glucose (filled circles). Speeds were measured in both aerobic (pink shaded area) and anaerobic (cyan shaded area) conditions. Gray shaded area gives the swimming speeds of cells prior to collection of AB30 (suspension A, squares) or of the cells harvested after $\approx 50\,\mathrm{min}$ of fermentation (suspension B, triangles).

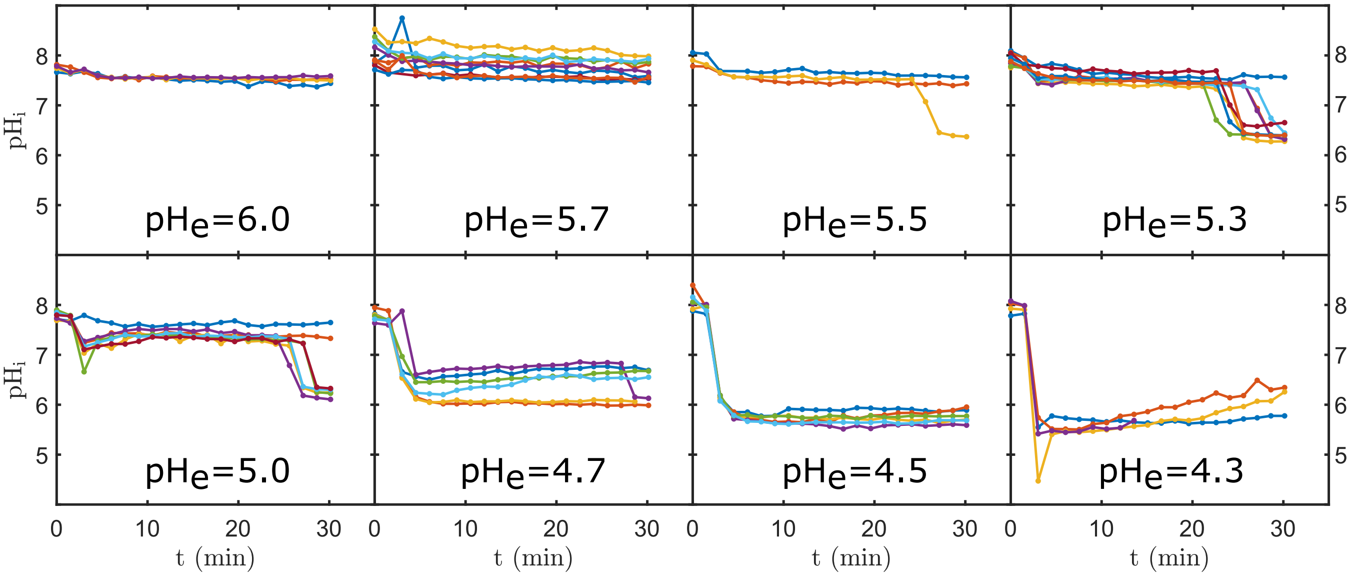


**Fig. SI13.** Internal pH measurements conducted at various $\mathrm{pH_e}$. Cells are initially in $1\times$ MB + $20\,\mathrm{mM}$ glucose with $\mathrm{pH_e} = 7.0$ ($t \leq 2\,\mathrm{min}$), then $1\times$ MB + $20\,\mathrm{mM}$ glucose containing various amounts of lactic acid (see Fig. 4) is flown into the tunnel slide (*Methods*). Each curve corresponds to the average value of the $\mathrm{pH_i}$ measured on 20-40 individual cells in the tunnel slide. Error bars are similar to those in Fig. 4(a) and are omitted for clarity.

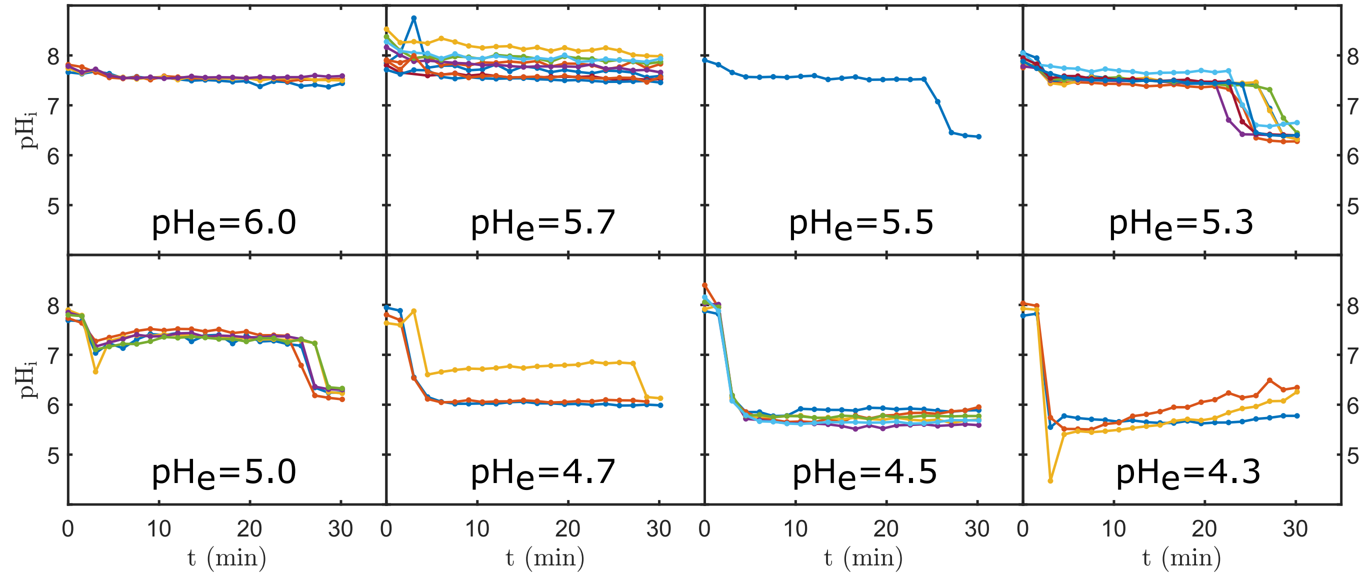


**Fig. SI14.** Internal pH measurements conducted at various $\mathrm{pH_e}$, displaying only the batches of cells deemed to have turned anaerobic during the measurements, as indicated by the sharp $\mathrm{pH_i}$ drop. The steep drop is consistent with the swimming speed drop, which was previously directly correlated with oxygen deprivation by measuring the fluorescence lifetime of a dye sensitive to dissolved oxygen [5]. Data taken from Fig. SI13.

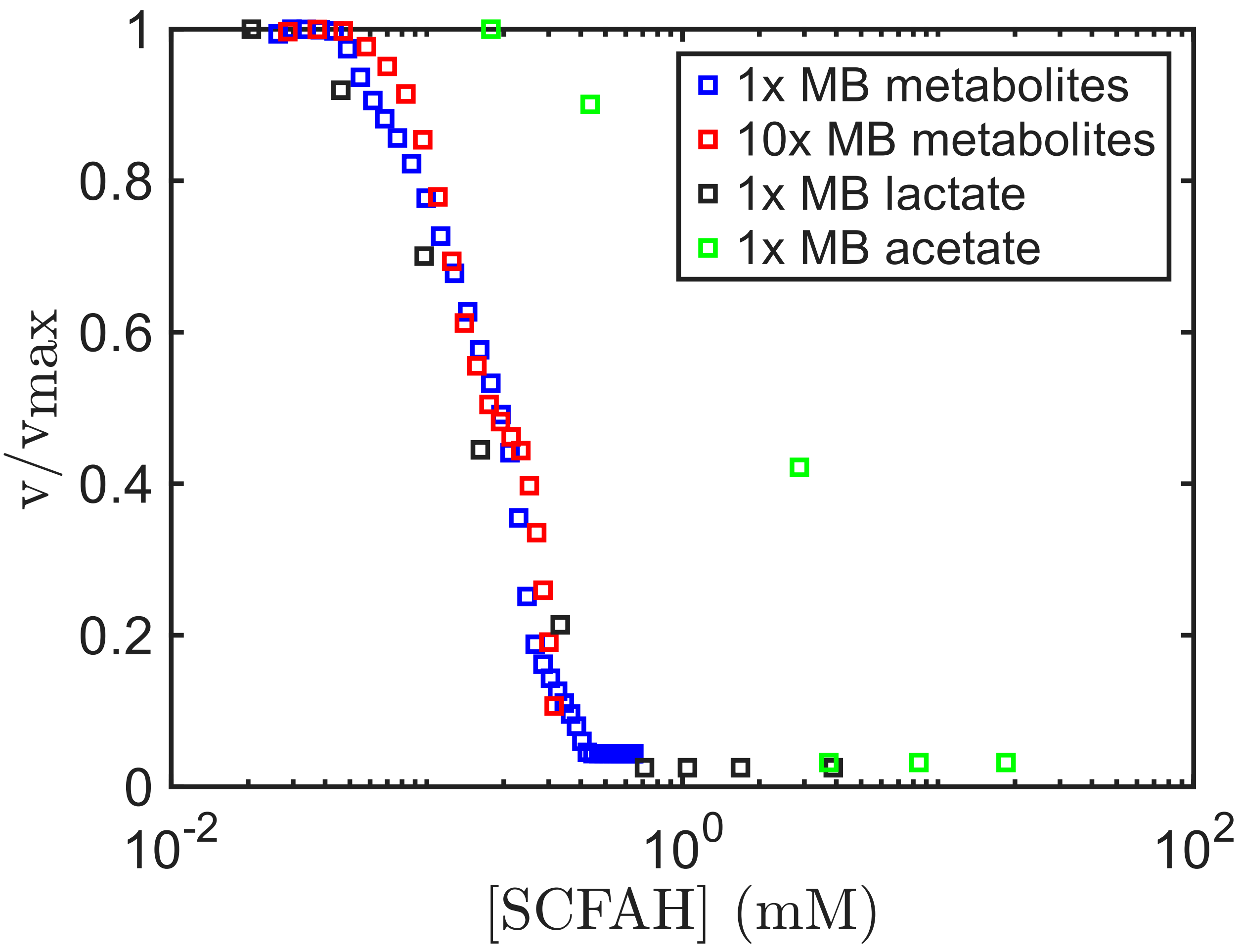


**Fig. SI15.** Comparison of the relationship between normalised speed and [SCFAH] for different SCFA under anaerobic conditions. A significantly larger concentration of protonated acetic acid (green squares) is required to cause a decay similar to that caused by lactic acid (black squares) or by the metabolites mixtures produced by cells during fermentation (blue and red squares).

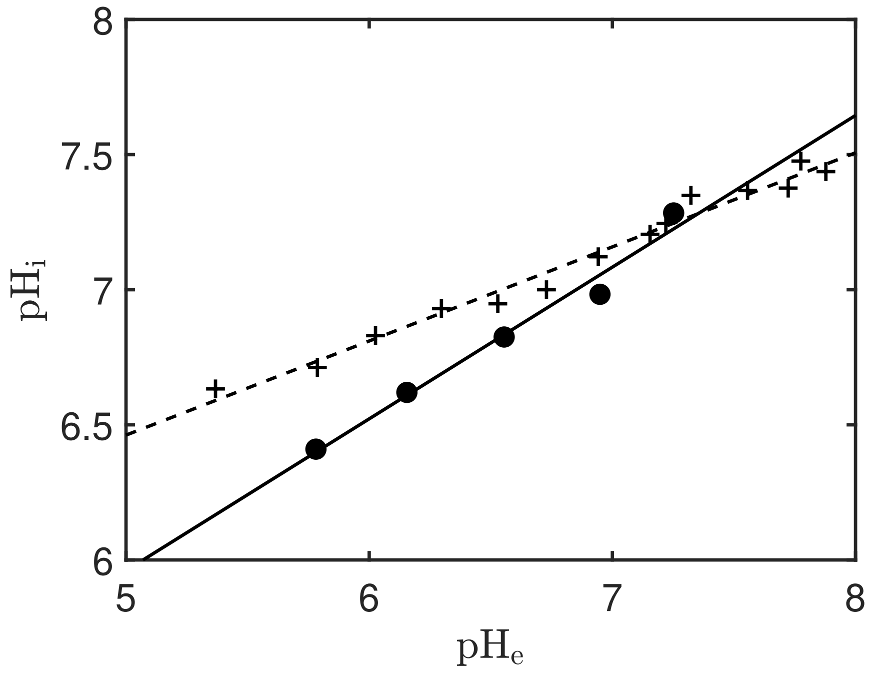


**Fig. SI16.** Manually extracted data from Ref. [6], with $\mathrm{pH_i}$ plotted against $\mathrm{pH_e}$ in MB + $10\,\mathrm{mM}$ lactate (crosses) and MB + $10\,\mathrm{mM}$ lactate + $34\,\mathrm{mM}$ acetate (circles). Black solid and dashed lines are best fit lines for each conditions: $\mathrm{pH_i} = 0.3483\mathrm{pH_e} + 4.7205$ for MB + $10\,\mathrm{mM}$ lactate, $\mathrm{pH_i} = 0.5615\mathrm{pH_e} + 3.1534$ for MB + $10\,\mathrm{mM}$ lactate + $34\,\mathrm{mM}$ acetate.
